# Embryonic dopaminergic neuron activity sustains lifelong locomotion in *Drosophila*

**DOI:** 10.64898/2025.11.28.691140

**Authors:** Aishwarya Padmanabhan, Daniel Rahman, Ruibao Zhu, Salah Khorbtli, Sarugan Sathianathan, Bastien Le Flohic, Lauren Assanga Bisse, Bertrand Mollereau, Cheng Huang, Nikos Konstantinides, Abdul Raouf Issa

## Abstract

Locomotor skills arise early in life and are maintained throughout an animal’s lifespan, yet how this continuity is achieved despite major neural remodeling remains unclear. Using *Drosophila*, which undergoes complete metamorphosis, we show that the activity of embryonically established dopamine neurons (DANs) is essential for locomotion across all developmental stages and adulthood. Through stage-specific behavioral assays, optogenetics, *in vivo* brain imaging, and fluorescent neuronal tracking, we identify a subset of ventral nervous system (VNS) DANs that modulate locomotor function throughout life. Transcriptomic analyses reveal that they maintain expression of developmental transcription factors. Knocking down these factors, particularly Antp and Pdm2, in post-mitotic VNS DANs reduces neurite arborization and impairs adult locomotion. These findings uncover a previously overlooked function for embryonic DANs and suggest that stable locomotion during nervous system maturation relies on persistent developmental regulator expression coupled with structural remodeling.

## INTRODUCTION

Locomotion is an evolutionarily conserved, innate motor activity that is executed throughout the lifespan. In many animals, it emerges early during development, often *in utero* or *in ovo*, where it appears first as primitive movements^1–4^. As development progresses, these primitive movements mature into more complex locomotor behaviors, including crawling and walking^1,4,5^. It is believed that the primitive movements rely on the activity of a set of nerve cells established in the spinal cord^2,6–9^, including glial cells^3^and sensory neurons^2,10^. Despite advances in identifying these early locomotor circuits, few studies have systematically investigated the relationship between the neurons driving primitive movements and those involved in locomotor behaviors later in life^1,11^. This remains a significant gap in our understanding of how locomotion is maintained across developmental stages despite ongoing structural and functional changes in the nervous system that affect neuronal anatomy, cell count, and transcriptomic profiles.

Dopaminergic neurons (DANs) are key modulators of locomotion across species^12–15^, and have received increased interest since the discovery of the role of dopamine in Parkinson’s disease (PD)^16^. In mammals, ascending midbrain DANs initiate locomotion, while descending DANs projecting to the brainstem and spinal cord mostly regulate the speed^12,13,17^. However, whether and how common locomotor DANs act from embryonic development through adulthood remain unclear. Given that DAN dysfunction underlies neurological disorders with motor components^18,19^ identifying the role of DANs from development to adulthood could provide new insights into the mechanisms underlying these diseases.

The fruit fly *Drosophila melanogaster* is an attractive model for studying the role of DAN in the maintenance of locomotion across life-stages. It has well-defined developmental stages, an accessible nervous system and powerful genetic tools that enable spatiotemporal manipulation of neuronal activity, structural investigation, and precise control of gene expressions^2,20–22^. *Drosophila* undergoes metamorphosis, transitioning from a larval to an adult form through a pupal stage^23^. During this process, the nervous system is extensively remodeled, some neurons are pruned or die, while others survive and refine their connections^24^. Among the surviving neurons are those of the locomotor system, including DANs, which are born during embryogenesis and whose activity is required to regulate locomotor behaviors such as walking, climbing, and flying^25–27^.

In this study, we showed that DAN activity is essential at the embryonic stage for proper locomotor performance later in life and in adulthood in flies. Using a combination of immunolabelling assays, optogenetics and reporter immortalization techniques, we uncovered a subset of DANs in the fly subesophageal zone (SEZ) and ventral nerve cord (VNC)—hereafter collectively referred to as the ventral nervous system (VNS)^28^—that are generated during embryonic development and persist through all stages of life to modulate locomotion. Furthermore, we showed through fluorescent labelling of neuronal membranes and RNAi-mediated gene knockdown, that the maintenance of VNS DAN function from development to adulthood relies on structural remodeling and the sustained expression of several developmental transcription factors (TFs), including the evolutionary conserved homeodomain genes such as Antennapedia (Antp), POU domain protein 2 (Pdm2) and Tup/Islet.

Our findings reveal a functional role for DANs during embryonic stages, establishing a link between early developmental processes and adult locomotion. They indicate that the maintenance of this function requires the stability of DAN cell body positioning and dopamine pathway gene expression, along with structural remodeling and continuous activity of developmental gene programs.

## RESULTS

### Embryonic suppression of DAN activity compromises locomotor performance later in life

Movement is present at all stages of Drosophila development, including peristaltic waves (hereafter referred to as twitching) at the embryonic stage^2,29^, crawling in larvae^14,30^, and walking in adults^5,31^. DANs and their receptors are present early during development, at the embryonic stage (**Figure 1A**), allowing us to test whether early DAN activity is essential for locomotor function development and, if so, to determine the consequences of chronic DAN inactivation during early development on locomotor maintenance later in life. To address these questions, we expressed the inward rectifying potassium channel Kir2.1, driven by the Tyrosine Hydroxylase line (TH-Gal4)^32^, from the embryonic stage (**Figure 1B**). Kir2.1 is commonly employed in the field to constitutively hyperpolarize neurons and suppress electrical activity^33,34^. The behavioral assay was performed using DeepLabCut^35^ to track larval body parts during crawling. This analysis revealed that chronic Kir2.1 expression alters larval locomotor patterns and performance, resulting in shorter travel distances and slower crawling speeds **(Figure 1C** and **Figure S1A**). Shibire^ts1^ (*shi*^ts1^), a temperature-sensitive mutation of the *Drosophila* gene encoding a Dynamin orthologue, blocks vesicle endocytosis and thus synaptic transmission and neurotransmitter release, at restrictive (elevated) temperature^36^. Similar to the effects observed with Kir2.1, larvae expressing *shi*^ts1^ in DANs and raised at the restrictive temperature (31°C) from the embryonic stage exhibit reduced locomotor performance (**Figure 1D** and **Figure S1B**). During normal development, late-stage larvae (L3) exhibit long-distance migration away from their hatching site to search for suitable conditions for feeding, growth, and pupation, a behavior known as wandering^37^. We observed that L3 expressing Kir2.1 in DANs failed to travel far, remaining near their hatching site or on the food medium (**Figure S1C**).

**Figure 1.**
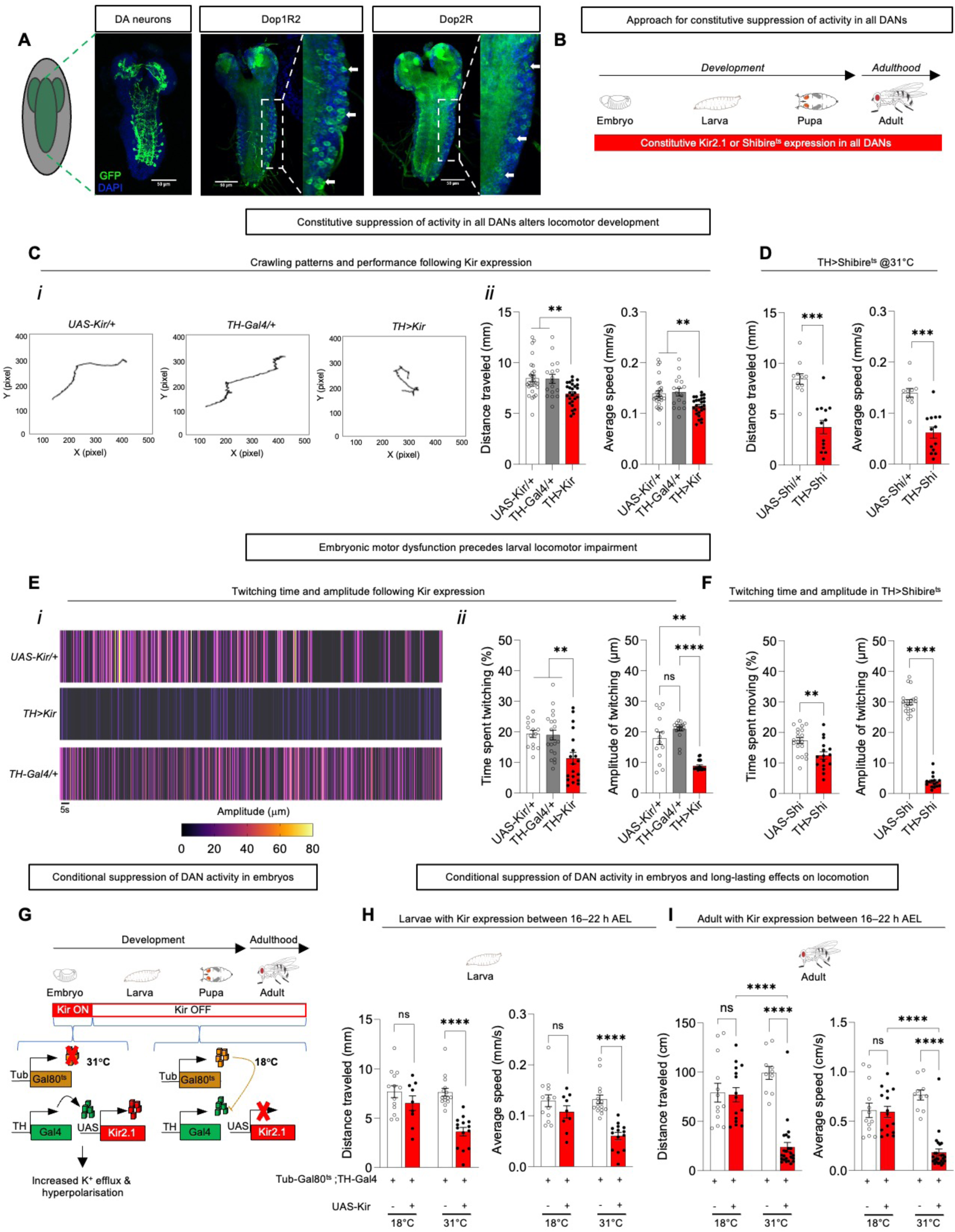
Suppression of DAN activity during embryogenesis disrupts locomotor development. **A.** Confocal images showing the presence of dopamine neurons (DANs) and dopamine receptors in the embryonic fly nervous system. They were visualized driving the Myr-GFP with TH-Gal4^32^ and *Dop1R2*-Gal4 and *Dop2R*-Gal4^96^ respectively. All scale bars are 50 μm. **B.** Approach for a constitutive suppression of activity in all dopamine neurons (DANs). Kir2.1^33^ or temperature-sensitive Shibire^TS36^ (at 31°C) was expressed in all DANs from the embryonic stage and across developmental stages. **C**. Constitutive suppression of activity in all DANs with Kir2.1^33^ (*TH>Kir*) alters locomotor development. A representative crawling pattern of the TH>Kir larva shows a reduced paths compared to controls (*UAS-Kir/+* and *TH-Gal4/+*) (i). This results in reduced distance traveled and lower crawling speed (ii). Data are represented as mean ± SEM. Error bars: 95% confidence interval (CI). ∗∗p < 0.01 by Mann-Whitney test. n for *UAS-Kir/+, TH-Gal4/+ and TH>Kir*: 28, 26 and 18, respectively. Data were collected from two independent experiments, and each animal was recorded individually. **D**. Constitutive suppression of activity in all DANs with Shibire^ts^ (*TH*>*Shibire^ts^*) alters locomotor development. Shibire^ts^ expressing larvae (*TH*>*Shi*), raised at 31°C during the embryonic stage, exhibit reduced locomotor speed and travel distance compared to controls (*UAS-Shi/+)*. Data are represented as mean ± SEM. Error bars: 95% CI. ∗∗∗p < 0.001 by Mann-Whitney test. n for *UAS-Shi/+ and TH>Shi*: 12 and 13, respectively. Data were collected from two independent experiments, and each animal was recorded individually. **E, F**. Altered locomotor behavior in larvae following constitutive expression of Kir2.1 (*TH*>*Kir*) or Shibire^ts1^ (*TH*>*Shibire^ts^*) in all DANs is preceded by impaired embryonic movements. Representative twitching patterns over time of the *TH>Kir* embryo and controls (*UAS-Kir/+* and *TH-Gal4/+*) (Ei). Heat hatches denote twitching. Both the timing and amplitude of embryonic twitching are reduced compared to controls (*UAS-Kir/+* and *TH-Gal4/+*) (Eii). Data are represented as mean ± SEM. Error bars: 95% CI. ∗∗p < 0.01 and ∗∗∗∗p < 0.0001 by Mann-Whitney test. n for *UAS-Kir/+, TH-Gal4/+* and *TH>Kir*: 14, 20 and 19, respectively. Each animal was recorded individually. A similar reduction in twitching timing and amplitude is observed in TH>Shibire^ts^ embryos (F). Data are represented as mean ± SEM. Error bars: 95% CI. ∗∗p < 0.01 and ∗∗∗∗p < 0.0001 by Mann-Whitney test. n for *UAS-Shi/+ and TH>Shi*: 20 and 16, respectively. Data were collected from two independent experiments, and each animal was recorded individually. **G**. Approach to suppress activity in all DANs exclusively during the embryonic stage. Kir2.1^33^ expression in all DANs is restricted to the embryonic stage. Targeted suppression of DAN activity specifically during embryogenesis was achieved using the temperature-sensitive, tub-Gal80^ts38^. At the restrictive temperature (31°C), Gal80^ts^ is inactivated, allowing *TH-Gal4*–driven Kir2.1 expression, leading to hyperpolarization of DANs via increased K⁺ efflux. At the permissive temperature (18°C), Gal80^ts^ represses Kir expression. **H, I**. Long-lasting locomotor effects following temporal suppression of activity in all DANs during the embryonic stage. Larvae that experienced Kir expression only during embryogenesis show reduced crawling distance and speed (H). n for *Tub*-*Gal80* ^ts^;*TH-Gal4/+* and *Tub*-*Gal80*^ts^;*TH>Kir* at 18°C: 13 and 10, respectively. n for *Tub*-*Gal80* ^ts^;*TH-Gal4/+* and *Tub*-*Gal80*^ts^;*TH>Kir* at 31°C: 15 and 14, respectively. Data were collected from two independent experiments, and each animal was recorded individually. Adults that expressed Kir2.1 only during embryogenesis also show reduced locomotor performance, including decreased distance traveled and walking speed (I). Data are represented as mean ± SEM. Error bars: 95% CI. (nonsignificant, ns) P>0.05 and ∗∗∗∗p < 0.0001 by Kruskal-Wallis test with Dunn’s multiple comparisons test n for *Tub-Gal80^ts^*;*TH-Gal4/+* and *Tub-Gal80*^ts^;*TH>Kir* at 18°C: 13 and 15, respectively. n for *Tub-Gal80^t^*^s^;*TH-Gal4/+* and *Tub-Gal80*^ts^;*TH>Kir* at 31°C: 10 and 25, respectively. Data were collected from two independent experiments, and each animal was recorded individually. See also Figure S1.

In light of the impaired larval locomotion, we examined whether embryonic movements were similarly affected in the absence of DAN activity. Using custom-made software (see STAR Methods) to assess movement in 17 h after egg laying (AEL) embryos expressing Kir2.1 or *shi*^ts1^, we observed a reduction in both the duration and amplitude of embryonic twitching (**Figure 1E,F, Figure S1D** and **Supplemental video 1**). These findings indicate that motor dysfunction arises during embryonic development, before the reduction in locomotion seen later in life.

The presence of motor dysfunction from the embryonic stage prompted us to assess the effect of activating DANs specifically during this period on the maintenance of locomotor behavior later in life. To provide supporting evidence to this, we suppressed DAN electrical activity selectively at the embryonic stage using the temperature-sensitive TARGET system, tub-Gal80^ts38^ (**Figure 1G)**. We noticed that suppressing activity specifically during 16-22h AEL, led to reduced travel distances and slower locomotor speeds in both larvae (**Figure 1H)** and adults (**Figure 1I)**. The 16-22h AEL window covers key developmental events, including the onset of TH protein expression^39^ and the emergence of spontaneous network activity^40^. As such, DAN activity during embryonic development is essential for the development of locomotion and its maintenance into adulthood. It also provides a model for studying how DANs affect locomotion from development to adulthood.

### Chronic suppression of DAN activity in the VNS impairs locomotion across development and adulthood

It is well established in flies that loss of structure in the adult dopaminergic system impairs locomotion^26,41^. This may therefore provide one possible explanation for the compromised locomotor function at later stages following embryonic suppression of DAN activity. To test this hypothesis, we performed immunolabelling of the larva L2 nervous system and observed a significant loss of neurite arborizations (**Figure 2A**), but not of cell bodies (**Figure S2A** and **S2B**), in DANs expressing Kir2.1. This reduction in arborization is particularly pronounced in the ventral nervous system (VNS)^28^, where our intersectional genetic strategy (otd-or Tsh-FLP combined with Tub>Gal80>^42^) restricting TH-Gal4 expression to the VNS revealed that most arborizations arise from local DANs (**Figure 2B** and **2C**). Therefore, we further explored the implication of VNS DANs in the locomotor alterations observed upon global pan-DAN activity suppression. To this end, we constitutively suppressed activity in DANs using the intersectional genetic strategy to restrict TH-Gal4-driven Kir2.1 expression to the VNS (**Figure 2D** and **Figure S2C**). Targeted suppression of VNS DANs resulted in locomotor alterations, observed as reduced embryonic twitching (**Figure 2E** and **Figure S2D**) and slower larval crawling (**Figure 2F** and **Figure S2E**), confirming the previously reported developmental role of spinal dopaminergic activity in shaping motor circuit structure and locomotor behaviors^43,44^. In addition, individuals who survived to adulthood under these conditions, exhibited impaired walking behavior (**Figure 2G** and **Figure S2F**). To further examine the temporal requirement of VNS DAN activity, we suppressed this activity exclusively during the embryonic stage (16-22h AEL) using Gal80^ts^ timing (**Figure 2H**). As observed with pan-DAN silencing (**Figure 1H,I**), suppression of VNS DAN activity during embryonic stages altered both larval and adult locomotion (**Figure 2I,J** and **Figure S2G,H**). We tested the efficiency of the conditional approach by assessing the ability of Gal80^ts^ to repress TH-Gal4-driven Kir expression in VNS DANs. We observed that larvae expressing Gal80^ts^, reared from the embryonic stage at the permissive temperature (18°C), exhibited undetectable Kir-GFP fluorescence in the VNS DANs, whereas those kept at the restrictive temperature (31°C) showed clear Kir-GFP expression (**Figure S2I**), demonstrating that the Gal80^ts^-based approach enables temporal control of Kir expression in VNS DANs. Collectively, these results identify VNS DANs as a key population whose early embryonic activity is required for locomotor development and maintenance in larvae and adults, respectively.

**Figure 2.**
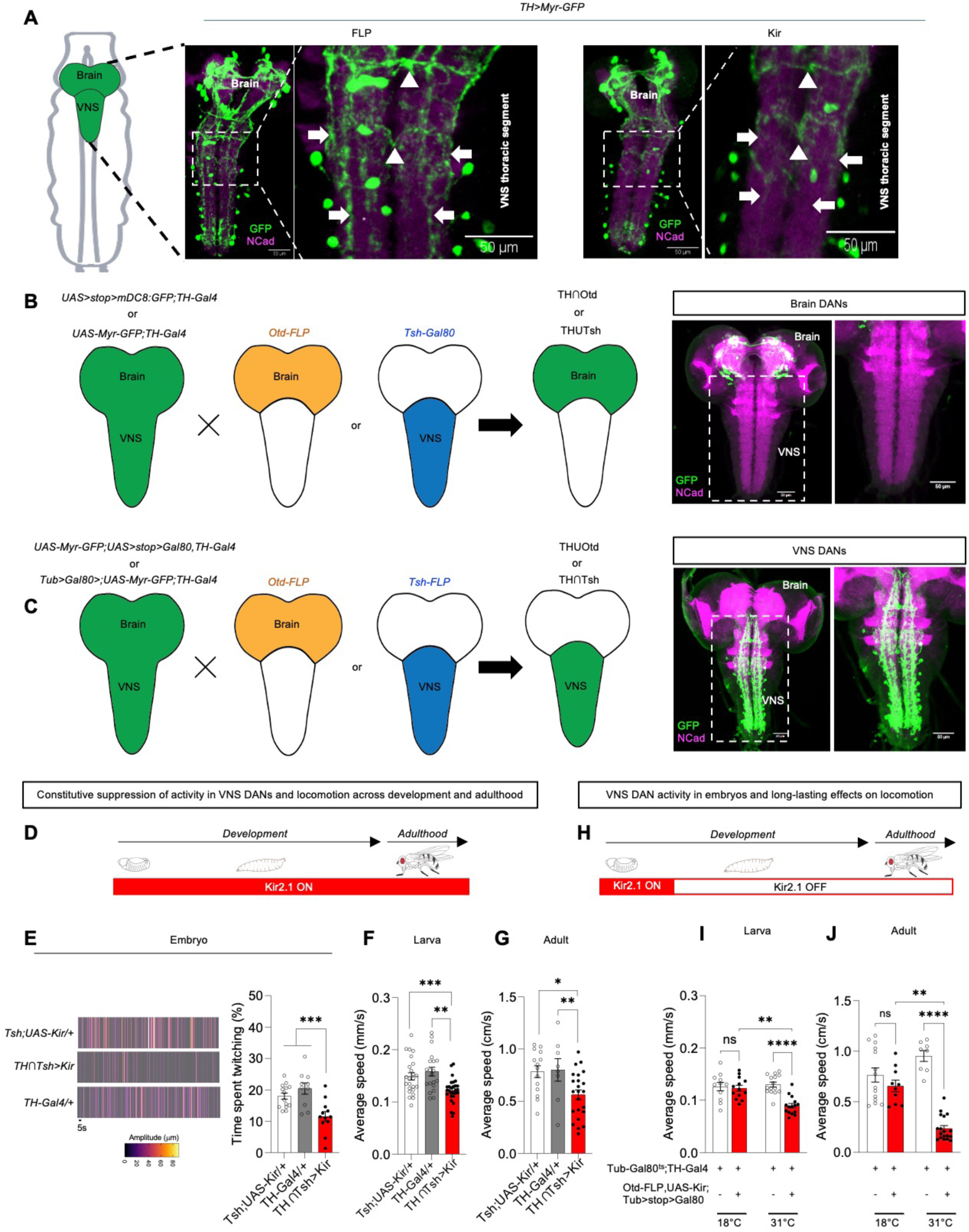
Suppression of DAN activity in the VNS is sufficient to impair locomotion across development and adulthood. **A**. A suppression of activity in DANs from embryonic stage reduces neurite arborizations in the larval L2 ventral nervous system (VNS). Representative confocal microscopy images of GFP-expressing neurite arborization of DANs within the VNS thoracic segment of larvae expressing Kir or FLP (control). Arrows and arrowheads indicate longitudinal and lateral projections, respectively. These decrease upon activity suppression (Kir expression), suggesting that both axonal and dendritic compartments may be affected. All scale bars are 50 μm. **B, C.** DAN neurite arborization pattern in the larval brain and VNS. Cartoon illustrating the intersectional genetic strategy used to specifically target brain DANs (B) (left): TH-Gal4 drives expression in both brain-and VNS-derived neurons (green). We crossed the TH-Gal4 line ‘*UAS>stop>mCD8:GFP;TH-Gal4*’ (green) with Otd-FLP (orange), which excises the stop cassette in the brain, or the TH-Gal4 line ‘*UAS-Myr-GFP;TH-Gal4*’ (green) with Tsh-Gal80 (blue), which represses Gal4 in the VNS. Both crosses can restrict TH-Gal4-mediated reporter GFP expression to the brain. Representative confocal images show that this intersectional genetic strategy enables labeling of brain DAN neurite structures with membrane-targeted myristoylated GFP (Myr-GFP) (B) (right). Cartoon illustrating the intersectional genetic strategy used to specifically target VNS DANs (C) (left): We crossed the TH-Gal4 line ‘*UAS-Myr-GFP;Tub>stop>Gal80,TH-Gal4*‘ (green) with Otd-FLP (orange), which excises the stop cassette in the brain, or the TH-Gal4 line ‘*Tub>Gal80>;UAS-Myr-GFP;TH-Gal4*’ (green) with Tsh-FLP (blue), which excises the stop cassette in the VNS. Both crosses restrict TH-Gal4-mediated reporter GFP expression to the VNS. Representative confocal images of larval L3 nervous system show that the intersectional genetic strategy enables labeling of VNS DAN neurite projections with membrane-targeted Myr-GFP (C) (right). Genotype: TH∩Tsh (*Tub>Gal80>; Tsh-LexA,FLP-LexAOP,UAS-Myr-GFP;TH-Gal4*). The VNS regions outlined by dashed boxes in (B, right) and (C, right) indicate that the majority of DAN neurite arborizations within the VNS originate from local neurons. All scale bars are 50 μm. **D-G**. Constitutive suppression of activity in VNS DANs alters locomotion during development and adulthood. Experimental approach involving Kir2.1^33^ expression in VNC DANs from the embryonic stage and across developmental stages (D). Expression of Kir2.1 in VNC DANs (TH∩Tsh>Kir) impairs embryonic movements (E). Representative twitching patterns over time of the TH∩Tsh >Kir embryo and controls (TshH;UAS-Kir/+ and TH-Gal4/+) (left). Heat hatches denote twitching. The time spent by the TH∩Tsh>Kir embryo twitching is reduced compared to controls (right). In the same experimental condition, TH∩Tsh*>*Kir larvae (F) and adults (G) exhibit slower speed. Data are represented as mean ± SEM. Error bars: 95% CI. ∗p < 0.05, ∗∗p < 0.01, and ∗∗∗p < 0.001 by Mann-Whitney test. n for UAS-Kir/+, TH-Gal4/+, and TH∩Tsh>Kir were 14, 10, and 12 (embryo); 24, 20, and 28 (larva); and 14, 8, and 23 (adult), respectively. Data were collected from two independent experiments, and each animal was recorded individually. **H-J**. Long-lasting locomotor effects following temporal suppression of activity in VNS DANs during the embryonic stage. Experimental approach involving Kir2.1^33^ exclusive expression in VNS DANs during the embryonic stage (H). Larvae (I) and adults (J) that have experience Kir expression at embryonic stage exhibit slower speed. Data are represented as mean ± SEM. Error bars: 95% CI. ∗p < 0.05, ∗∗p < 0.01, ∗∗∗p < 0.001 and ∗∗∗∗p < 0.0001 by Kruskal-Wallis test with Dunn’s multiple comparisons test. n for white and red plots at 18°C: 11 and 13 (larva); 10 and 14 (adult), respectively. n for white and red plots at 31°C: 15 and 15 (larva), 8 and 18 (adult) respectively. Data were collected from two independent experiments, and each animal was recorded individually. See also Figure S2.

### VNS DANs exhibit spatial and numerical stability across developmental life stages

The impact of chronic embryonic suppression of VNS DAN activity on locomotor maintenance in larvae and adults suggests that these neurons may also be present at throughout post-embryonic stages. To test this hypothesis, we mapped VNS DANs across developmental stages using TH-Gal4 to drive expression of a nuclear-localized GFP reporter (UAS-Stinger). Cell body distribution analyses revealed that VNS DANs occupy the same positions within the ventral medial and dorsal lateral regions of the thoracic and abdominal segments from embryo to adult (**Figure 3A-D**). Cell body counts showed that the number of DANs in the VNS remains constant across embryonic, larval, and adult stages (**Figure 3E**). Of note, stability of the VNS DAN number contrasts with the increased number of DANs seen in the brain, where new clusters such as PPM1/2/3 (protocerebral posterior medial (PPM3), and protocerebral anterior lateral (PAL) are observed in adults, new neurons are added within embryo-born clusters (**Figure 3E**). Given that the fly metamorphosis involves extensive neuronal cell death and reprogramming during the transition from larval to adult, and that adult nervous system originate from two waves of neurogenesis—embryonic and larval^24,45^, it is plausible that adult VNS DANs may not derive from the embryonic VNS DANs we characterized. In addition, while the TH-Gal4 driver was commonly used for the mapping of DANs at distinct developmental stages^46^(**Figure 3A**), its expression across all DANs from embryonic to adult stages has not been systematically verified. Therefore, to confirm the spatial and numerical stability of VNS DANs, we employed an immortalization strategy to trace these neurons across larval and adult stages of the nervous system. This approach includes a sequence of three genetically defined events (**Figure 3F**). Firstly, the temporally controlled heat inactivation of GAL80^ts^ to drive TH-GAL4 activity selectively at embryonic stages; secondly, the GAL4-induced Flippase-mediated recombination of stop codons flanked by FRT sites, which are located downstream of the polyubiquitin promoter (Ubip63) and upstream of the GFP gene; and thirdly, subsequent Ubip63-driven expression of GFP in neurons that once were TH-positive. Using this approach, we observed a comparable number and location of GFP-positive cells in both larval and adult SEZ, as well as within the thoracic and abdominal segments of the VNC (**Figure 3G,H** and **Figure S3A-C**). We conclude that VNS DANs persist from embryo to adult, preserving their spatial distribution and cell number.

**Figure 3.**
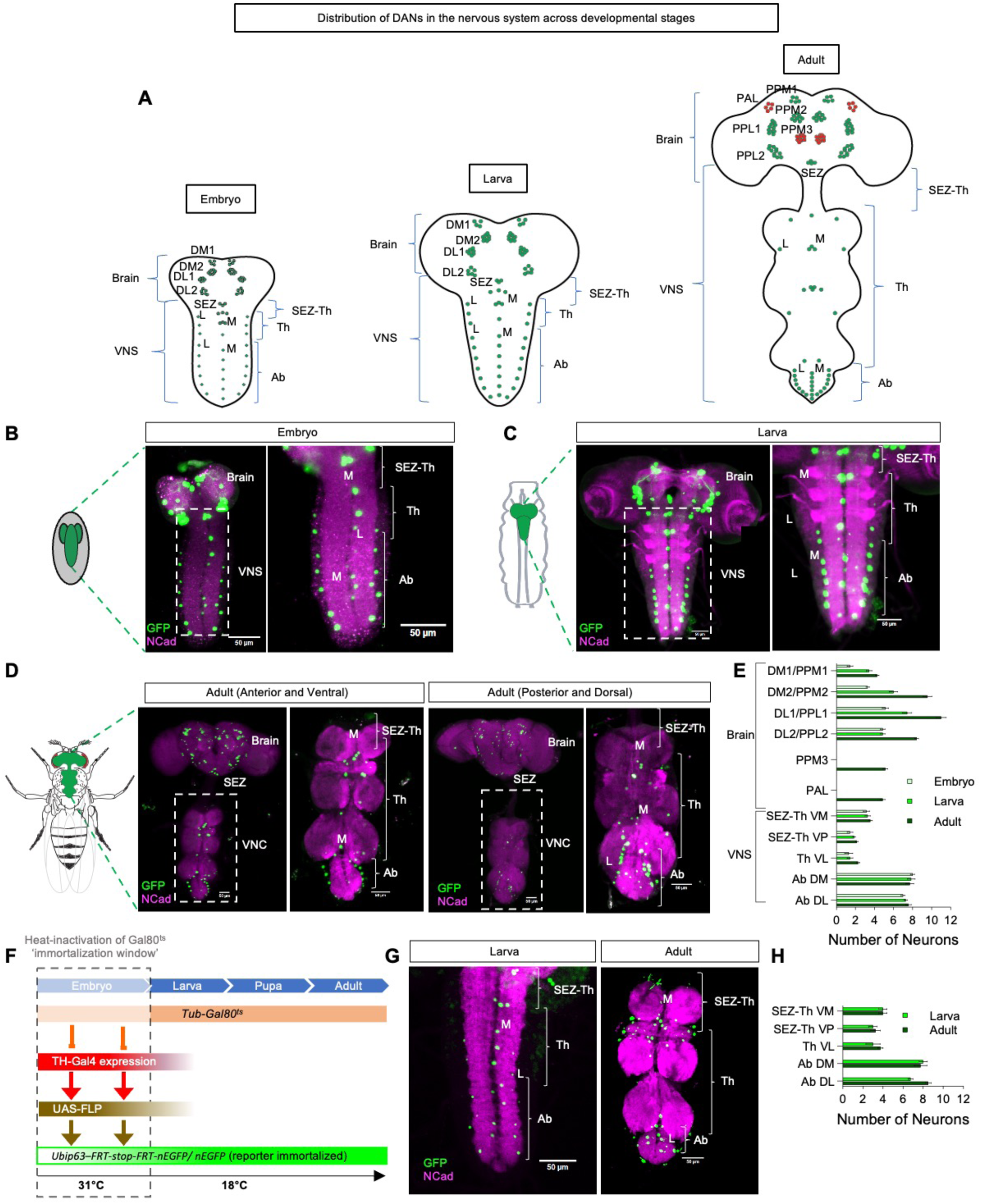
VNS DANs are spatially and numerically stable across developmental stages. **A**. Cartoon of the brain and VNS illustrating the previously documented nomenclature and number of DAN clusters mapped by TH-GAL4 at distinct developmental stages^39,46,74,107^. Embryo-born DAN clusters persist into the adult nervous system, which additionally contains the PAL and PPM3 clusters (highlighted in red). DL1, DL2, DM1, and DM2 clusters in embryos and larvae correspond to the adult DAN clusters PPL1, PPL2, PPM1, and PPM2, respectively. Abbreviations: DM1/2, dorso-medial clusters 1 and 2; DL1/2, dorso-lateral clusters 1 and 2; PPL1, protocerebral posterior lateral 1; PPM1/2/3, protocerebral posterior medial 1/2/3; PAL, protocerebral anterior lateral; Ab, abdominal lateral neurons; Th, thoracic; SEZ, subesophageal zone; L, lateral; M, medial. **B–E.** Spatial distribution and number of DANs across developmental stages. Confocal images of embryos (17h AEL) (B), larvae (L2–L3) (C), and adults (D) expressing nuclear GFP (Stinger) in DANs highlight the organization of SEZ-Th, Th, and Ab clusters. Quantification of DAN cluster numbers per hemisphere in embryos (17h AEL), larvae (L2–L3), and adults shows that VNS DANs remain numerically stable across life stages (E). n= 6-7 individual animals. All scale bars are 50 μm. **F–H**. Immortalization of a DAN reporter. Schematic of the approach (F). This strategy involves three genetically defined steps: (1) heat-induced inactivation of GAL80^ts^ to enable TH-GAL4 activity during the embryonic stage; (2) GAL4-driven Flippase-mediated recombination to remove FRT-flanked stop codons located between the polyubiquitin promoter (Ubip63) and nuclear GFP (nEGFP); and (3) subsequent Ubip63-driven nEGFP expression in neurons that were previously TH^+^ during the embryonic stage. Spatial distribution of nEGFP^+^ DANs in larval and adult VNS (G) reveals no difference in cell numbers. n=4 individual animals. See also Figure S3. All scale bars are 50 μm.

### Structural remodeling of VNS DANs across development is essential for adult locomotion

Embryo-born neurons that persist into the adult nervous system typically undergo during metamorphosis a dramatic structural remodeling that leads to changes of axon, dendrite and synapse architecture through mechanisms such as pruning and/or regrowth^24,47–49^. To investigate if VNS DANs undergo such remodeling, we employed membrane-tagged myristoylated GFP (Myr-GFP) to fluorescently label neurite arborizations, including dendrites and axons, across different developmental stages. We observed that VNS DAN neurites arborize differently in the VNC neuropil during embryonic and larval development compared to adulthood (**Figure 4A**-**C**). In adulthood, the VNS DAN neurite arborizations is pruned, with prominent axon projections visible in the brain neuropil, particularly in the superior medial protocerebrum, anterior midline, and periesophageal neuropils. This observation suggests that although VNS DANs persist with identical cell body positions and numbers across developmental stages, their arborization patterns are remodeled during nervous system maturation, a process that may support functional transition from development to adulthood. We argued that the distinct arborization patterns result from the remodeling process and support functional transition from development to adulthood. Consistent with this idea, we find that a disruption of the remodeling process by targeting cell adhesion or guidance effector genes, using RNA interferences (RNAis) against Beat-Ib, Beat-Ic and Toll-6^50,51^ (**Figure S3D**), which are enriched in the VNS DANs^22^ impairs locomotor performance (**Figure 4D**). We further observed that downregulation of these genes reduces DAN neurite arborization in the VNS during the larval-to-adult transition (**Figure 4E**, **F**), supporting their role in neuronal remodeling.

**Figure 4.**
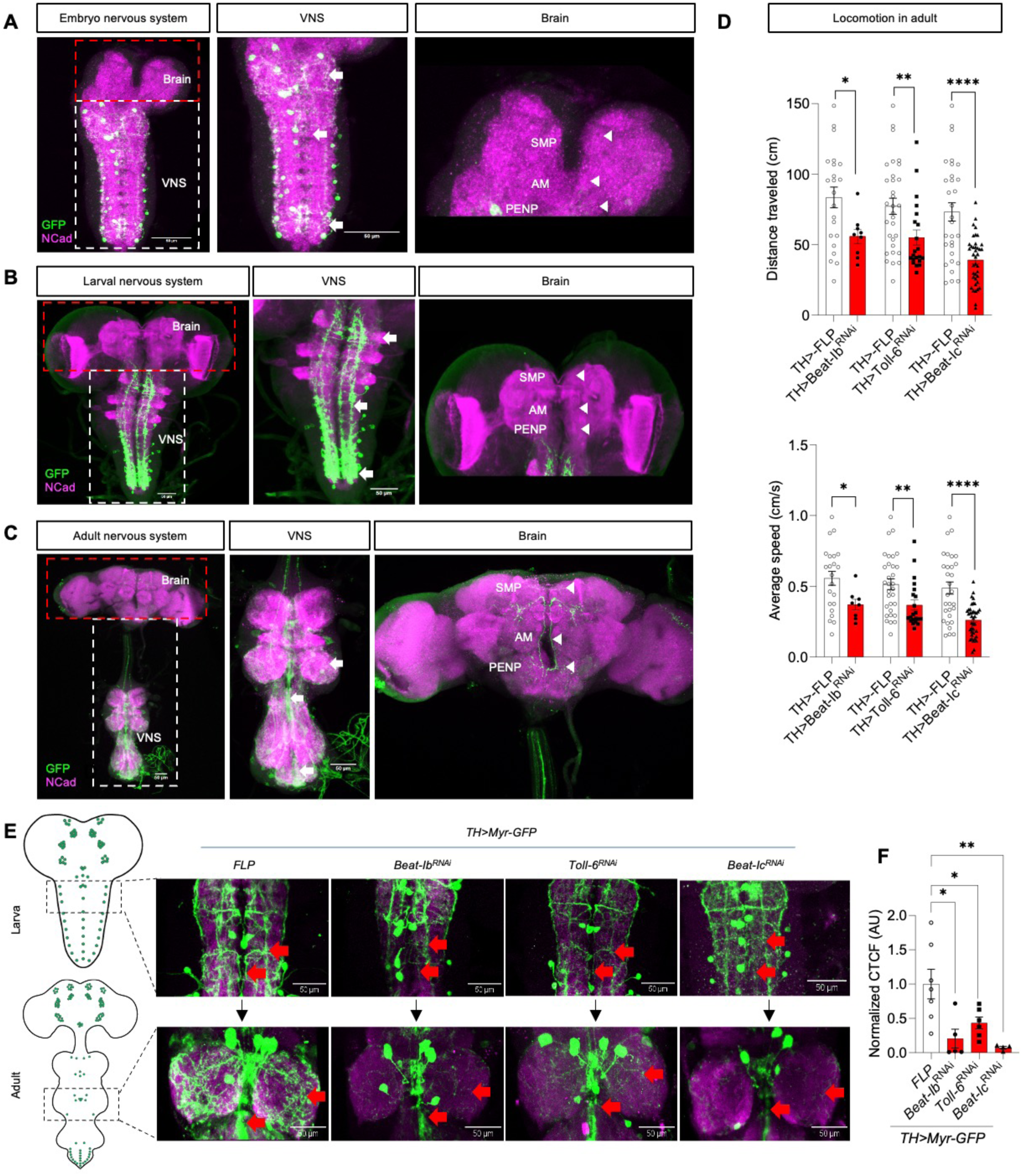
**Transition of VNS DANs from embryo to adult requires structural remodeling. A–C**. Structural remodeling of VNS DANs across distinct developmental stages. Confocal images of the nervous system in embryos (A) and larvae (B) show that VNS DANs exhibit dense neurite arborizations within the VNS (white arrows, white dashed boxes, and middle panels), with no projections into the brain (right panels). In adults (C), VNS DANs display pruned arborizations in the VNS (middle panel, dashed box) and extensive projections into the brain (right panel), which indicate structural remodeling of these neurons during the transition from embryo to adult. Arrows indicate longitudinal and lateral projections within VNC neuromeres, and arrowheads indicate a specific area in the brain targeted by mature projections. SMP: superior medial protocerebrum, AM: anterior midline, PENP: periesophageal neuropils. All scale bars are 50 μm. **D**. Disruption of the remodeling process impairs locomotion in adult. Downregulation of cell adhesion or guidance effector genes highly enriched in VNC DANs, leads to reduced distance traveled and decreased average locomotor speed in adults. Data are represented as mean ± SEM. Error bars: 95% CI. ∗p < 0.05, ∗∗p < 0.01, and ∗∗∗∗p < 0.0001 by Mann-Whitney tests. n for *TH>FLP* (22, 31, and 30), *TH>beat-Ib^RNAi^* (18), *TH>Toll-6^RNAi^* (21), and TH*>beat-Ic^RNAi^*. (30). Data were collected from two independent experiments, and each animal was recorded individually. **E**, **F**. Downregulation of cell adhesion or guidance effector genes affects DAN remodeling. Representative confocal images showing DAN neurite arborization within the VNC neuromeres of *TH>beat-Ib^RNAi^*, *TH>Toll-6^RNAi^* and TH*>beat-Ic^RNAi^* larvae and adults (E). All scale bars are 50 μm. Red arrows indicate longitudinal and lateral projections. Quantification of adult DAN arborization density, measured as corrected total cell fluorescence (CTCF) (F). These panels show a reduction in neurite arborization following beat-Ib, Toll-6 and beat-Ic RNAi-mediated knockdown. Data are represented as mean ± SEM. ∗p<0.05, and ∗∗p < 0.01 by Mann-Whitney tests. n=4-6. See also Figure S3.

### Acute activation of DANs in VNS during ongoing behavior modulates locomotor speed at distinct developmental stages

During locomotion, animals must continuously adjust speed and gait in response to changing internal states and environmental stimuli^52–54^. Such rapid adjustments often involve neuromodulators^55–58^. Therefore, having established that VNS DANs persist from embryonic to adult stages and that their early silencing impairs locomotor development and maintenance, we hypothesized that VNS DAN activity also plays a modulatory role during ongoing locomotion across developmental stages, consistent with the role of neuromodulators in controlling motor circuit excitability and output vigor^55–57,59,60^. To this end, we expressed the red-light-activated channelrhodopsin Chrimson::mVenus—which enables rapid, acute, and reversible neuronal activation^61^—in VNS DANs and assessed the effects of transient red-light stimulation on locomotor behavior at distinct developmental stages. In embryos, activation of VNS DANs significantly reduces the duration of twitching without affecting its amplitude (**Figure 5A, B** and **Supplemental video 2**). At later developmental stages, red-light stimulation decreases both larval crawling (**Figure 5C, D** and **Supplemental video 3** and **4**) and adult walking (**Figure 5E, F** and **Supplemental video 5** and **6**) respectively. Conversely, transient inactivation of VNS DANs using the green-light gated chloride channel GtACR1^62^ led to enhanced locomotor performance in embryos (**Figure S4A, B**), larvae (**Figure S4C, D**), and adults (**Figure S4E, F**). These optogenetic results indicate that VNS DAN activation negatively modulates the execution of locomotor behavior, indicating that a transient activation of these neurons is necessary to slow or halt locomotion. These results suggest that, in addition to their role in the development and maintenance of locomotion, as revealed by Kir2.1-mediated silencing (**Figure 2E-G, 2I** and **2J**), acute activation of VNS DANs exerts a dynamic or real-time modulatory influence on locomotor output.

**Figure 5.**
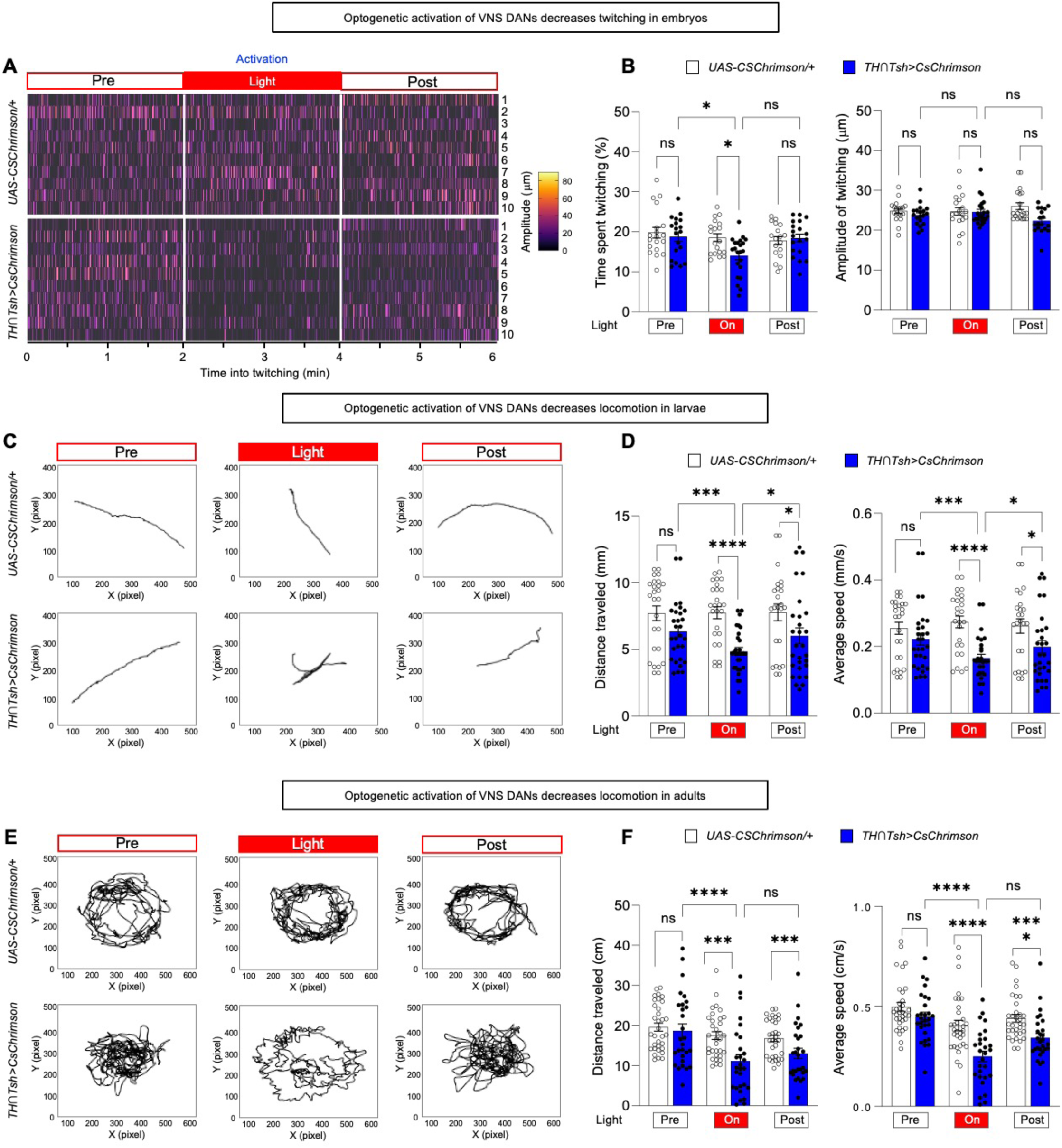
Acute activation of VNS DANs modulates locomotor speed across distinct developmental stages. **A, B**. Embryonic twitching. Heatmaps show that red-light stimulation decreases twitching in embryos expressing Chrimson::mVenus in VNC DANs (TH∩Tsh>CsChrimson), compared to pre-and post-light periods and genetic controls (UAS-CsChrimson/+) (A). Heat hatches denote twitching and each row represents individual embryo. Quantification of time spent twitching and twitching amplitude (B). Data are represented as mean ± SEM. Error bars: 95% CI. ns p>0.05 and ∗p < 0.05 by Mann Whitney tests for comparisons with parental controls; Friedman tests followed by Dunn’s tests for within-genotype comparisons. n for UAS-CsChrimson/+ and TH∩Tsh>CsChrimson for all periods: 18 and 19, respectively. Data were collected from two independent experiments, and each animal was recorded individually. **C, D**. Larval locomotion. Representative crawling patterns show shorter paths in TH∩Tsh>CsChrimson larvae during red-light stimulation, compared to pre-and post-light periods and controls (UAS-CsChrimson/+) (C). Quantification of distance traveled and crawling speed (D). Data are represented as mean ± SEM. Error bars: 95% CI. ∗p < 0.05, ∗∗∗p < 0.001 and ∗∗∗∗p < 0.0001 by Mann Whitney tests for comparisons with parental controls; Friedman tests followed by Dunn’s tests for within-genotype comparisons. n for UAS-*CsChrimson*/+ and *TH*∩*Tsh*>*CsChrimson* for all periods: 26 and 30, respectively. Data were collected from two independent experiments, and each animal was recorded individually. **E, F**. Adult locomotion. Representative walking patterns show shorter paths in *TH*∩*Tsh*>*CsChrimson* adults during red-light stimulation, compared to pre-and post-light periods and controls (UAS-CsChrimson/+) (E). Quantification of distance traveled and walking speed (F). Data are represented as mean ± SEM. Error bars: 95% CI. ns p > 0.05, ∗∗∗p < 0.001 and ∗∗∗∗p < 0.0001 by Mann Whitney tests for comparisons with parental controls; Friedman tests followed by Dunn’s tests for within-genotype comparisons. n for UAS-*CsChrimson*/+ and *TH*∩*Tsh*>*CsChrimson* for all periods: 26 and 24, respectively. Data were collected from two independent experiments, and each animal was recorded individually. See also Figure S4.

When a DAN is activated, it releases dopamine into the synaptic cleft. The released dopamine then binds to dopamine receptors (DopRs) on postsynaptic neurons, triggering intracellular signaling cascades that ultimately influence behavior ^12,63^. If activation of VNS DANs reduces locomotor performance, then targeted activation of neurons expressing DopRs should elicit similar effects. We therefore optogenetically activated neurons expressing the two major Drosophila dopamine receptors, D1-and D2-like, using Dop1R2-and Dop2R-Gal4 lines^64^. Activation of Dop1R2⁺ neurons, but not Dop2R⁺ neurons, led to a strong reduction in locomotor performance across developmental stages, in embryo (**Figure 6A** and **Figure S5A**), larva (**Figure 6B** and **Figure S5B**), and adult (**Figure 6C** and **Figure S5C**), further confirming that acute dopaminergic activity reduces locomotion. Our Dop1R2⁺-Gal4 expression mapping (**Figure S5D**) shows that Dop1R2⁺ neurons are broadly distributed across the fly brain and VNS, suggesting that the observed locomotor phenotype may arise from the activation of heterogeneous neuronal populations^65,66^.

**Figure 6.**
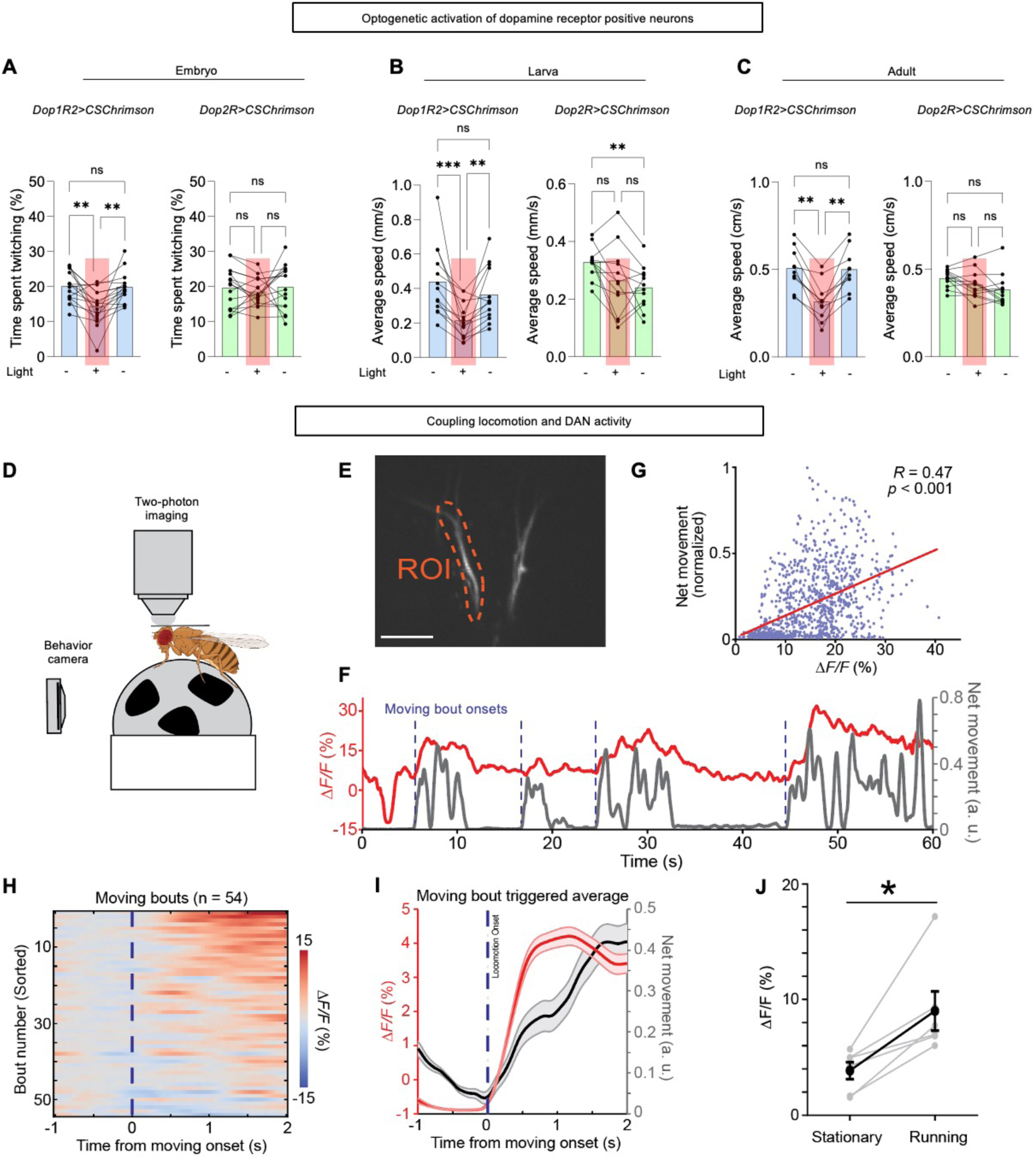
Acute activity of VNS DANs is correlated with locomotor performance. **A–C**. Activation of dopamine receptor–positive neurons correlate with the locomotor activity. Activation of Dop1R2^+^ neurons reduce locomotion at distinct developmental stages. Red-light stimulation decreases embryonic twitching (A), larval crawling (B), and adult walking (C) in animals expressing Chrimson::mVenus in Dop1R2^+^ neurons (Dop1R2>CsChrimson) (left), compared to pre-and post-light periods and Dop2R>CsChrimson flies (right). Data are represented as mean ± SEM. ns p>0.05, ∗p < 0.05, ∗∗p < 0.01 and ∗∗∗p < 0.001 by one-way ANOVA with Dunnett’s T3 multiple comparisons test. n for Dop1R2>CsChrimson and Dop2R>CsChrimson for all periods: 16 and 16 (embryo), 13 and 12 (larva), and 10 and 12 (adult). Data were collected from two independent experiments, and each animal was recorded individually. **D–F**. Neural activity of VNC DANs correlates with fly locomotion. Schematic of the experimental setup. A tethered fly with a surgical window in its cuticle walks on an air-suspended ball during *in vivo* two-photon functional imaging (D). A behavioral tracking camera simultaneously monitors the ball’s rotation for calculating the fly’s moving trajectory. Representative image of GCaMP8s expression in axons of the VNC DANs within the central brain (E). The dashed line outlines the region of interest (ROI) used for extracting calcium activity. Scale bar 20 µm. Example traces showing calcium activity from the axons of the VNC DANs (red) and the fly’s corresponding net movement (gray) during a 60-second imaging session (F). a.u., arbitrary units. Blue dashed lines indicate the onsets of moving bouts that have sustained locomotion (> 2s) following a short rest (>1s). Correlation between the neural activity and net movement from the traces in panel F (G). The best-fit line (red), Pearson correlation coefficient (R), and P-value indicate a statistically significant positive relationship (*P* = 1×10^-58^). Heatmap of neural activity in the ascending axons of the VNC DANs aligned to the onset of sustained locomotion (>2s) following a short rest (>1s) (H). Each row represents a single movement bout. The 54 bouts, collected from 6 flies, are sorted by average calcium activity during the 0–2 s moving phase. The vertical blue line marks the onset of movement. Mean traces of neural activity (red) and locomotion (gray) averaged from the 54 movement bouts shown in panel H (I). Shaded areas represent the SEM. The vertical blue line marks the onset of movement. Comparison of average calcium activity during stationary (net locomotion < 0.05 a.u. for ≥ 1 s) and running (net locomotion > 0.15 a.u. for ≥ 1 s) periods (J). Gray dots represent individual flies, and the black dots and line represent the mean. ∗p<0.05 by Wilcoxon signed-rank test. n = 6 animals, recorded individually. See also Figure S5.

To further understand the relationship between the activation of VNS DANs and decreased locomotor performance, we monitored VNS DAN activity from axonal projections within the central brain during adult walking via two-photon calcium imaging (**Figure 6D**). GCaMP8s signals were recorded from VNS DAN axons in the brain of tethered flies walking on an air-supported ball (**Figure 6D, E**). We observed that elevated GCaMP8 signals correlate with episodes of accelerated walking or leg movements (**Figure 6F, G** and **Supplemental video 7**), indicating that VNS DAN activation occurs during locomotor bouts. Interestingly, as the fly transitions from rest to running, the initial increases in GCaMP/calcium signals and moving speed are nearly simultaneous. However, the calcium signal demonstrates a slower ramping dynamic thereafter compared to the increased moving speed (**Figure 6H–J**). Together, our optogenetic and *in vivo* calcium imaging experiments demonstrate that VNS DAN activity dynamically modulates basal locomotor speed during ongoing behavior.

### Persistence of developmental genetic programs is required in VNS DANs for lifelong locomotor maintenance

In addition to the gene coding for tyrosine hydroxylase (TH), the rate-limiting enzyme in dopamine synthesis, VNS DANs also express key dopamine pathway genes, including the vesicular monoamine transporter (VMAT) and the dopamine transporter (DAT), throughout development and into adulthood^22,67^ (**Figure S6A, B**). This sustained expression suggests the presence of a regulatory program that controls the expression of dopamine pathway genes across VNS DAN lifespan. To test this hypothesis, we performed a cross-analysis of available single-cell transcriptomic datasets^22,68^ to compare the expression of transcription factors found in embryonic and adult VNS DANs (**Figure S6A, B**). We found that several transcription factors (TFs)—hereafter referred to as developmental TFs—traditionally described for their roles in early steps of neural development, such as neural progenitor proliferation and neuronal differentiation^69^, remain expressed either broadly in neurons or specifically in DANs of the adult VNS (**Figure 7A**, **Figure S6A, B** and **Supplemental Table S1**). Among these, we identified key evolutionarily conserved homeodomain developmental TFs such as Antp/Hox6-8, a POU domain protein homologous to Oct-2 (Pdm2/Oct2), and a LIM-HD homolog (Tup/Islet)^69^(**Figure 7A**). Given that these TFs are known for their roles during neural differentiation^69^, we next tested whether their persistent expression in post-mitotic neurons, such as VNS DANs, is required for function. To this end, we used different RNAis (**Figure S6C**) to constitutively knockdown (KD) Antp/Hox6-8, Pdm2/Oct2, and Tup/Islet and assessed effects on adult locomotion. Our experiment showed that KD of Antp, Tup, or Pdm2 resulted in reduced adult locomotor performance (**Figure 7B** and **Figure S6D**). Importantly, we found that conditional knockdown of Antp and Pdm2 specifically in adult DANs also impairs locomotion (**Figure 7C,D** and **Figure S6E**). Given the known developmental role of homeoproteins in regulating neuronal identity^69^ and the link between structural integrity and locomotion^26,41^, we examined DAN morphology upon Antp or Pdm2 knockdown. Strikingly, we observed a reduction in neurite arborization within the VNC neuromeres (**Figure 7E,F**). We further confirmed that Antp and Pdm2 are translated into protein in both embryonic and adult VNS DANs (**Figure S6F**). Collectively, these results indicate that the locomotor defects may involve structural alterations caused by the loss of these transcription factors. They also suggest that the expression of developmental TFs is part of intrinsic molecular programs that contribute to VNS DAN function and the regulation of locomotion.

**Figure 7.**
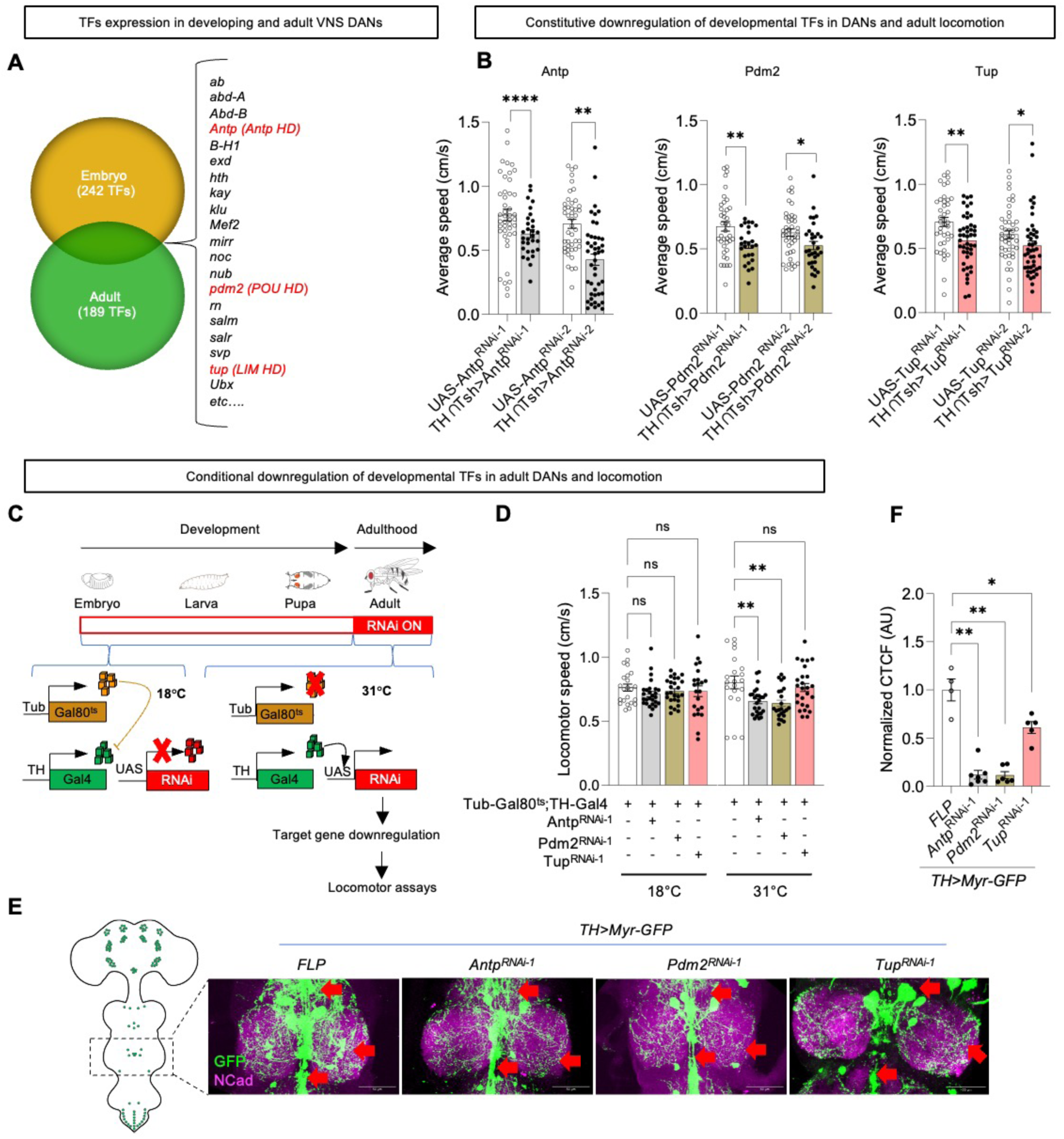
Developmental genetic programs are reused by VNC DANs to sustain locomotor function. **A.** Two hundred transcription factors were found to be expressed both in embryonic and adult dopaminergic neurons. Twenty of them are displayed in this Figure. The rest can be found in Supplemental Table S1. Three of them (in red) were selected for behavioural experiments. **B.** Constitutive RNAi-mediated downregulation of selected developmental transcription factors, including Antp, Pdm2, and Tup, impairs adult locomotion. Data are represented as mean ± SEM. Error bars: 95% CI. ∗p < 0.05, ∗∗p < 0.01, and ∗∗∗∗p < 0.0001 by Mann-Whitney tests. n for *UAS-Antp^RNAi-1^*, *TH∩Tsh>Antp^RNAi-1^*, *UAS-Antp^RNAi-2^*, *TH∩Tsh>Antp^RNAi-2^*, *UAS-Pdm2^RNAi-1^*, *TH∩Tsh>Pdm2^RNAi-1^*, *UAS-Pdm2^RNAi-2^*, *TH∩Tsh>Pdm2^RNAi-2^*,*UAS-Tup^RNAi-1^*, *TH∩Tsh>Tup^RNAi-1^*, *UAS-Tup^RNAi-2^* and *TH∩Tsh>Tup^RNAi-2^*: 45, 30, 44, 45, 39, 26, 42, 32, 41, 45, 45, and 45. Data were collected from three independent experiments, and each animal was recorded individually. **C**-**F**. Downregulation of selected developmental transcription factors specifically in adult DANs impairs locomotion and neuronal morphology. Conditional strategy to exclusively downregulate Antp, Pdm2, and Tup in adult DANs (C). At the permissive temperature (18°C), Gal80^ts^ is active and represses Gal4-mediated gene (RNAi) expression. After adult eclosion, flies were shifted to the restrictive temperature (31°C), allowing Gal80^ts^ inactivation and Gal4-mediated RNAi expression (‘ON’). Locomotor assays were conducted after 10 days to ensure maximal degradation of transcription factor proteins translated prior to Gal80^ts^ inactivation. Expression of RNAi against Antp and Pdm2 in DANs impairs locomotor maintenance in adult flies (D). Data are represented as mean ± SEM. ns p >0.05 and ∗∗p < 0.01 by Kruskal-Wallis test with Dunn’s multiple comparisons test. n for *Tub-Gal80^ts^;TH-Gal4/+*, *Antp^RNAi-1^*, *Pdm2^RNAi-1^* and *Tup^RNAi-1^*transgenic lines were 23, 27, 26, and 22 at 18°C, and 22, 21, 26, and 23 at 31°C, respectively. Data were collected from three independent experiments, and each animal was recorded individually. Representative confocal images of DAN neurite arborization in adult VNS mesothoracic (E). All scale bars are 50 μm. Red arrows indicate longitudinal and lateral projections. Quantification of adult DAN arborization density, measured as corrected total cell fluorescence (CTCF) show that downregulation of Antp, Pdm2 or Tup in adult DANs reduce neurite arborization (F). Data are represented as mean ± SEM. ∗p<0.05, and ∗∗p < 0.01 by Mann-Whitney tests. n=5. See Figure S6.

## DISCUSSION

Activity of specific neuronal subsets in the nervous system, particularly within the spinal cord, has been shown to influence the formation of motor circuits during embryonic development and contributes to the early acquisition of locomotor skills^2,8,9^. However, nervous systems undergo significant changes in neuronal number, morphology, and function after birth or hatching and throughout life^45,70–72^. How the neuronal subsets that drive early locomotion persist through or adapt to these developmental transitions remains poorly understood. Addressing this question is important to understanding the onset of human neurodevelopmental and neurodegenerative disorders, many of which involve dopaminergic or locomotor dysfunction^18,19^. One explanation for this gap has been experimental tracking of individual neurons within the locomotor system from embryo to adult. Using the *Drosophila* locomotor system, we show that embryonically active VNS DANs bridge early-stage movements to adult locomotor performance. The functional continuity of these DANs is associated with substantial structural remodeling coupled with preservation of a stable transcriptional identity.

### Embryonic DAN activity is essential for locomotor development and maintenance

Dopamine neurons (DANs) are considered the key modulator of locomotor behaviors across species^12–15^. In this study, we observe that chronic embryonic suppression of activity in DANs alters locomotor development and results in long-lasting impairments of locomotion in adults. This finding indicates that dopaminergic locomotor function emerges during embryogenesis and that this emergence sets the stage for the establishment of adult motor circuits. While DANs have been anatomically described in the fly embryo^39,73,74^, our study is the first to demonstrate their functional relevance at this developmental stage. This observation aligns with studies in vertebrates that showed that DANs^75–78^, as well as populations of interneurons, sensory neurons, and motor neurons^7^ exhibit synaptic activity, early during development, demonstrating that they become functionally active well before adulthood. In humans and rodents, motor function spans the entire lifespan, with its maturation during late embryonic and early postnatal periods being critical for the acquisition and maintenance locomotor behavior^1,79^, as developmental dysregulation of dopaminergic signaling impact function, with sustained effects on the physiology and behavior that can persist throughout adulthood^80^. As such, DANs are neuronal components supporting the locomotor continuity between embryonic and adult forms of organisms or species.

### Acute activation of VNS DANs during ongoing behavior modulates locomotor speed

We found that neuronal subsets from VNS (VNS DANs) are a part of DANs whose acute activation during locomotion reduces speed in both developing and adult flies. The VNS includes SEZ-VNC, can be considered analogous to the vertebrate brainstem and spinal cord, following the identification of the midbrain–hindbrain boundary (MHB) as a significant organizing center underscoring an evolutionarily conserved subdivision of the nervous system in both vertebrates and flies^81,82^. In this framework, the anterior region of the MHB corresponds to the protobrain, while the posterior region corresponds to the protocord^81,82^. Accordingly, VNS DANs may have already been present in the limbed bilaterian common ancestor, suggesting that these neuronal subsets in flies and vertebrates share conserved anatomical and functional features. Nonetheless, recent studies have shown that the VNS hosts many neurons whose activity controls behaviors such as sleep and motivation^83,84^, indicating that this region of the nervous system is not just a simple equivalent of the mammalian spinal cord receiving commands from the brain, but rather a control center for multiple neuronal functions.

In limbed animals, neuronal circuits that coordinate locomotor behaviors are located in the brainstem and spinal cord regions and established early during development^9,13,85^. These regions convey sensory information, receive commands from the brain, and send appropriate motor output instructions to the peripheral. Therefore, the fly VNS is expected to harbor neurons whose activity controls movements of locomotor apparatus and modulates the speed of the locomotion at all stages of development. It has been described that DANs of brainstem modulate locomotion in vertebrates^12,13,86^. Importantly, recent work has shown that hindbrain and spinal cord DANs reduce locomotor activity^12^, highlighting a potential evolutionary parallel with VNS DAN function in fly locomotion.

More recently, in the fly, optogenetic stimulation of DANs in flies via TH-Gal4 decreases locomotor activity^87–89^, whereas selective activation of protocerebral anterior medial (PAM) neurons using R58E02-Gal4 leads to increased locomotion^87^. These findings support the idea that some DAN groups including VNS DANs help slow down movement in flies. Consistent with this, we also observed a strong inhibitory effect on locomotion following the activation of Dop1R2⁺ neurons. In mammals, locomotion is primarily modulated by two populations of dopamine-sensitive neurons within brainstem and basal ganglia circuits: neurons expressing D1 receptors promote locomotion by reducing inhibitory output, whereas neurons expressing D2 receptors exert the opposite effect^12,63^. In *Drosophila*, homologs of D1-and D2-like receptors are Dop1R2 and Dop2R, respectively^90,91^. Based on this analogy, activation of Dop1R2⁺ neurons are expected to enhance locomotion. One possible explanation for the opposite effect observed here is that Dop1R2 may differ functionally from canonical D1-like receptors in *Drosophila*^92–95^, suggesting that it may have distinct roles in invertebrate locomotor control. Another possibility is that Dop1R2 is broadly expressed across the fly nervous system, being present in multiple neuronal populations, as suggested by our transgenic expression mapping (**Figure S5D**) and single-cell transcriptomic datasets^65,66^, indicating that Dop1R2⁺ neurons might comprise a heterogeneous population including glutamatergic, GABAergic, and cholinergic neurons. In addition, axon labeling of Dop1R2 neurons showed that these neurons project to the motor center (AMMC)^96^. Thus, activation of Dop1R2⁺ neurons likely recruits distinct neurotransmitter-defined cell types with both inhibitory and excitatory functions. In *Drosophila*, motor neurons are glutamatergic and form excitatory synapses onto muscles, whose activation induce sustained muscle contraction and thereby inhibits locomotion^97–99^. These motor neurons may be included in the Dop1R2⁺ neuronal population. Interestingly, a large-scale activation screen of the Janelia GAL4 collection reported locomotor phenotypes resembling those observed following activation of VNC DANs or Dop1R2⁺ neurons^100^. In that study, activation of specific neuronal populations in larvae slowed crawling through tonic contraction, whereas others displayed repeated ventral contractions followed by attempts to crawl. Future studies combining Dop1R2 and neurotransmitter-defined intersectional lines with connectomic analyses will be required to determine whether these specific neuronal populations express the Dop1R2 receptor and form anatomical and functional connections with VNC DANs. Taken together the effects of acute activation of VNS DANs and Dop1R2⁺ neurons across embryonic, larval, and adult stages, we propose a model of dopaminergic modulation of locomotor behavior that extends beyond ascending brain pathways across the animal kingdom. In this model, dopamine signaling via VNS DANs functions as a modulator or ‘brake’ on locomotion, counterbalancing the acceleration driven by other neuronal inputs^14,101,102^.

### Persistence and remodeling of DANs in the VNS support locomotor maintenance

Most neurons in the adult nervous system are post-mitotic and do not undergo significant replacement, meaning that many neurons born in their early life-stages persist for the entire lifespan without replacement^103–106^. This requirement is especially striking in flies due to metamorphosis^24,47^. In this study, we demonstrate a spatial and numerical stability of DANs in the VNS across distinct developmental stages. Our observations are consistent with the work of Hartenstein and colleagues, who reported that all clusters of embryonic-born DANs survive metamorphosis and persist into the adult fly VNS^46,107^. Neurons that survive metamorphosis and persist across the fly developmental stages undergo structural remodeling^24,108^. The remodeling process and functional importance that we documented for VNC DAN echoes this process and underscores its evolutionary conservation from invertebrates to vertebrates. Indeed, remodeling in vertebrate locomotor circuits was first documented by Ramón y Cajal, who described “process resorption” of spinal motor neurons, where early arborizations are pruned and reorganized into mature patterns^109^. Since then, structural remodeling has been recognized across species as a universal mechanism essential for the development, maturation and adaptability of neural circuits throughout life^49,108,110,111^. Accordingly, impairments in this process can manifest as alterations in adult locomotor performance, as observed in our study when interfering with molecular mediators of remodeling in DANs, or as the basis of neurological disorders^112–114^.

### Expression of developmental regulators in VNS DANs maintain locomotor function

In many organisms, the persistence of neurons for the entire lifespan without renewal suggests that the genetic systems that are required for supporting neuronal function must be maintained over years or as long as the animal lives^115^. We show that embryonic and adult VNS DANs share transcription factor expression, which is essential for maintaining locomotor function. Transcriptomic and GWAS evidence have reported such continuous expression of developmental TFs in mature nervous systems^22,116,117^, and such expression has been shown to regulate various aspects in mature neurons, including axonal targeting^118^, neurotransmitter identity^119,120^, and neuronal activity^121^. In mice, adult midbrain DANs maintain the expression of TFs such as *Pitx3, En1* and *Nurr1,* that are initiated during development, to preserve their dopaminergic identity and functional properties^122–124^. Similarly, long-term function of developmental TFs including Hox proteins and terminal selector *ast-1* and *ceh-43* are reported in the worm^119,125^. Recently, our study from *Drosophila* confirmed the continuous function of developmental TFs in the mature post-mitotic neurons by showing that a downregulation of the Hox protein *Ubx* in DANs leads to reduced neuronal activity necessary for flight locomotion^121^. Among the TFs expressed in both embryonic and adult VNS DANs, we identified Antp, Pdm2, and Tup, and found that the continuous expression of Antp and Pdm2 in adult DANs is particularly required for locomotor function and neurite arborization maintenance. These transcription factors have been previously described in neuronal differentiation, as well as in various aspects of postmitotic neurons, including survival, dendritic morphology, and neurotransmitter identity during development^69,119,126,127^. For example, Tup has been shown to regulate axon pathfinding and neurotransmitter identity in the fly DANs^73^. As such, these findings support a pervasive role for developmental regulators in controlling specific aspects of neuronal identity that support locomotor function after development has concluded. Future studies targeting additional determinants of dopaminergic identity and using knockout strategies, will be necessary to determine molecular pathways that Antp or Pdm2 to locomotor function maintenance in adulthood.

In this study, we identify, for the first time, the functional role of DANs during the embryonic stage, establishing a link between early neural activity and the maintenance of locomotor function in adulthood. Our findings reveal that a subset of DANs preserves key features of its cellular identity, including cell body position, morphology, functional properties, and the expression of terminal effector genes such as those involved in cell adhesion and guidance, as well as dopamine synthesis or transport pathway (e.g., pale, VMAT and DAT). Despite this stability, these neurons undergo substantial structural remodeling across developmental stages while maintaining the expression of several developmental transcription factors from embryo to adult. Together, these findings suggest that lifelong maintenance of locomotor function depends on a finely tuned balance between neuronal identity stability and structural plasticity, and that this process may be regulated by the persistent expression of developmental gene regulators.

### Limitations of the study

We show that RNAi-mediated knockdown of cell adhesion and guidance genes (*beat-Ib*, *Toll-6*, *Beat-Ic*) and developmental transcription factors (*Antp*, *Pdm2*, *Tup*) reduces neurite arborization in adulthood. However, whether *beat-Ib*, *Toll-6*, and *Beat-Ic* are direct or indirect transcriptional targets of *Antp* and/or *Pdm2* remains untested. In addition, although dopaminergic identity and function depend on dopamine synthesis and transport machinery, expression of key genes such as *TH*, *VMAT*, and *DAT* was not assessed upon knockdown of *Antp*, *Pdm2*, or *Tup*. This limits mechanistic interpretation of how these factors contribute to DAN function maintenance. Finally, structural analyses under activity suppression or transcription factor knockdown did not determine which neurite compartments (dendritic, axonal, and/or synaptic) are differentially affected. Therefore, future studies linking *Antp* and or *Pdm2* regulation, neuronal activity, and dopamine pathway and cell adhesion/guidance programs will be essential to define the mechanisms underlying lifelong maintenance of embryonically specified DANs.

## Lead contact

General inquiries should be addressed to and fulfilled by the lead contact, Abdul Raouf Issa.

## Materials availability

Any additional information, including Drosophila strains used in this study, is available from the lead contact.

## Data and code availability

- Custom-made code generated for this study has been deposited in Zenodo under distinct DOIs listed in the STAR Methods section and is publicly available as of the date of publication.
- Single cell mRNA sequencing data were downloaded from Gene Expression Omnibus (GSE141807 and GSE202987) and analyzed using code available on GitHub via https://doi.org/10.5281/zenodo.20125919.
- Any additional information required to reanalyze the data including raw behavioral and microscopy generated in this paper is available from the lead contact upon request.

## ACKNOWLEDGMENTS

We thank S. Birman, M. Crickmore, M. Zlatic, the Bloomington Stock Center, and the Vienna Drosophila RNAi Center for fly stocks, as well as the Brain Plasticity Laboratory fly food service collaborators, C. Beauchamp and A. Didelet. We are also grateful to S. Birman, T. Preat, M. Crickmore, F. Besse, and D. Rogulja and P. Faure for their valuable comments on an earlier version of this work. We acknowledge Ludovic Olanier for 3D printing the behavioral arenas. We also thank the Marie Skłodowska-Curie Actions program N°101067877 to Raouf Issa in Bertrand Mollereau laboratory, whose funding contributes to collect preliminary data for this project. This work was supported by grants from the French Labex Memolife, ATIP-Avenir, and ANR ERC-Tremplin programs, as well as by Université PSL’s Major Research Programs.

## AUTHOR CONTRIBUTIONS

R.I. designed the study. R.I. and A.P. performed all the experiments and data analyses with inputs of D.R. except for *in vivo* brain imaging experiments and transcriptomic dataset analyses, which were performed by C.H. and N.K. respectively. R.Z. prepared flies and contributed to imaging analysis codes, S.K. developed the behavioral setup and FicFrac analysis pipeline. R.I., B.L., S.S. and L.A-B. developed behavioral analysis software and code. G.W., T.M. and N.A. conducted behavioral experiments. R.I. wrote the manuscript with input from B.M., A.P., C.H. and N.K.

## DECLARATION OF GENERATIVE AI AND AI-ASSISTED TECHNOLOGIES IN THE WRITING PROCESS

During the preparation of this manuscript, the author(s) used ChatGPT to refine and streamline portions of the text. All content was subsequently reviewed and edited by the author(s), who take full responsibility for the content of the publication.

## SUPPLEMENTAL INFORMATION

**Document S1. Figures S1–S6**

---

**Supplemental video 1. Motor performance of embryos expressing Kir in DANs, Related to Figure 1**. Kir expression in DANs reduces the time the embryo spends moving.

**Supplemental video 2. Motor performance of embryos upon red-light stimulation, Related to Figure 5**. Red-light stimulation does not affect the time into twitching of control embryos (UAS-CsChrimson, top). In contrast, red-light stimulation reduces twitching duration in embryos expressing CsChrimson in VNS DANs (TH∩Tsh>CsChrimson, bottom). Related to Figure 5.

**Supplemental video 3. Locomotor performance of control larvae (UAS-CsChrimson) upon red-light stimulation, Related to Figure 5**. Red-light stimulation does not affect crawling speed in UAS-CsChrimson larvae.

**Supplemental video 4. Locomotor performance of larvae expressing CsChrimson in VNS DANs (TH∩Tsh>CsChrimson) upon red-light stimulation, Related to Figure 5**. Red-light stimulation reduces crawling speed in TH∩Tsh>CsChrimson larvae.

**Supplemental video 5. Locomotor performance of control adult flies (UAS-CsChrimson) upon red-light stimulation, Related to Figure 5**. Red-light stimulation does not affect walking speed in UAS-CsChrimson adult flies.

**Supplemental video 6. Locomotor performance of adult flies expressing CsChrimson in VNS dopaminergic neurons (TH∩Tsh>CsChrimson) upon red-light stimulation, related to Figure 5**. Red-light stimulation reduces walking speed in adults expressing CsChrimson in VNS DANs.

**Supplemental video 7. Two-photon calcium imaging in a behaving fly, Related to Figure 6**. Top left, ΔF/F movie showing GCaMP8s fluorescence signals from axonal terminals of VNC DANs in the central brain. Middle left, video from camera 1 capturing the front view of the fly running on an air-suspended trackball beneath the microscope objective. Bottom left, video from camera 2 providing a lateral view of the same behavior. Right, simultaneously recorded calcium activity (blue) from VNC DAN axons and the fly’s net movement (red) over a 60-s imaging session. a.u., arbitrary units. Videos displayed at 5× real time.

**Table S1**. List of transcription factors that are expressed in both adult and embryo VNS DANs, Related to Figure 7 and Figure S6.

**Table S2**. List of primer sequences used for qPCR analysis, Related to Figure S3 and Figure S6.

## EXPERIMENTAL MODEL AND STUDY PARTICIPANT DETAILS

### Fly husbandry and strains

Fruit flies, *Drosophila melanogaster* were maintained at 25°C in standard culture tubes containing water, cornflour, yeast, agar, methyl 4-hydroxybenzoate in a temperature-controlled incubator at 50% humidity with a 12 h/12 h cycle of alternating light and dark. Both male and female flies were used equally for the behavioral, imaging, and qPCR experiments, except in embryonic assays, where we were unable to distinguish sex. Unless otherwise indicated experiments were conducted under room temperature. For temperature switch manipulations flies were reared at 18°C, or 25°C, or 31°C at the time stages indicated.

The following Gal4 drivers were used: TH-Gal4^32^ (as kindly provided by Serge Birman, CNRS, ESPCI Paris Tech, France); Dop2R-Gal4; Dop1R2-Gal4^96^ (gift from Yi Rao). Effector and reporter lines included UAS-Kir:eGFP^128^ (RRID:BDSC_6595 and 6596); UAS-myr:GFP^129^ (RRID:BDSC_32197); UAS-CSChrimson^61^ (RRID:BDSC_55136), UAS-GtACR1^62^ (RRID:BDSC_92983); UAS-stinger (RRID:BDSC_84277); UAS-Shibire^ts^ (RRID:BDSC_44222); UAS-IVS-RSET-jGCaMP8s (RRID:BDSC_605078 and 605079); UAS-FLP (RRID:BDSC_4539); and Ubi-p63E(FRT.STOP)Stinger (RRID:BDSC_32250).

Additional transgenic lines *otd*-*FLP*,tub>*stop>Gal80* (Gift from Mike Crickmore) and Tub>Gal80>; Tsh-LexA,LexAOP-FLP (Gift from Marta Zlatic) were employed. We also used RNAi lines UAS-Toll-6RNAi (RRID:BDSC_64968); UAS-Beat-IcRNAi (RRID:BDSC_64528); UAS-Beat-IbRNAi (RRID:BDSC_55938); UAS-AntpRNAi-1 (RRID:VDRC_KK101774)^130^; UAS-AntpRNAi-2 (RRID:BDSC_27675); UAS-Pdm2RNAi-1 (RRID:BDSC_50665); UAS-Pdm2RNAi-2 (RRID:VDRC_GD52272); and UAS-TupRNAi-1 (RRID:VDRC_KK103585), and UAS-TupRNAi-2 (RRID:BDSC_51763). Finally, Tub-Gal80^ts^ lines (RRID:BDSC_7018 and 7019) were employed for temporal experiments, and w^1118^ was crossed to parental lines for controls.

## METHOD DETAILS

### Behavioral experiments

#### Locomotor behavior analysis

We analyse locomotor behaviors in embryos, larvae, and adults by recording contractions, crawling and walking, respectively.

Embryos: Parental flies were maintained in small collection cages supplied with apple juice agar plates supplemented with yeast paste. Embryos were collected after 18 hours, and stage 17 embryos were selected, according to their phenotypes for contraction recordings between 17–21 hours after egg laying (AEL), following the previous procedure^131^. Embryos were dechorionated using double-sided Scotch tape and transferred into custom 3D-printed chambers containing 3 × 3 mm wells. The wells were coated with dissolved Scotch tape glue in heptane, and embryos were covered with a layer of halocarbon oil (50:50 mixture of Sigma Halocarbon oil 27 H8773-100ML and 700 H8898-50ML) to prevent desiccation during recording. Recordings were performed using a Leica Ivesta 3 stereomicroscope (C-mount) equipped with a Basler ACE acA1300-200um monochrome USB 3.0 camera. The entire setup was placed inside a Comgrow protective enclosure (750 × 700 × 900 mm) to maintain stable ambient temperature and humidity. Videos were acquired at 30 frames per second (fps) and analyzed using a combination of Fiji (ImageJ) and custom software named Lifelong Movement Analysis Software (LIMAS). This software delineates the morphology and contour of the embryo and generates an ellipsoidal region of interest (ROI) around it. The variation in the ROI area (ΔA) according to the calculation described below. A= π × L/2 × W/2, with L and w embryo length and width respectively.

LIMAS outputs were exported as csv files, which were then processed further using a custom-made Python code (also available at https://doi.org/10.5281/zenodo.20074821) to determine the time the embryo spent contracting and the amplitude.

Larvae: Stage 17 embryos were selected and transferred to a fresh plate for 36 h. Freshly hatched larvae (L2) were placed on 0.9% agarose plates and were allowed to acclimatize for 2 minutes. To record forward locomotion (crawling) freely moving larvae were recorded for 2 minutes at a rate of 30 fps with a Leica Ivesta 3 stereomicroscope (C-mount) equipped with a Basler ACE acA1300-200um monochrome USB 3.0 camera. Videos were analysed with the open source software DeepLabCut^35^ (see method below) and the total travelled distance, speed and velocity quantified using a custom-made Python code (also available at https://doi.org/10.5281/zenodo.20074985).

Adults: 7-10 days individual adult fly was placed into a 3D-printed acrylic arena (2×2×0.5cm) with sloped edges^132^, covered by a transparent plexiglass (1mm thick). The freely walking fly is recorded for 5 minutes at a rate of 30 fps with a Basler ACE acA1300-200um monochrome USB 3.0 camera and TECHSPEC® 6mm UC Series Fixed Focal Length Lens. Videos were analysed with the open source software DeepLabCut^35^ (see method below) and the total travelled distance, speed and velocity quantified using a custom-made Python code (also available at https://doi.org/10.5281/zenodo.20074769).

#### Optogenetic assays

Embryos: Parental flies were reared in the dark in the vials containing retinal food (regular fly food mixed with 1mM all-trans retinal (Sigma)) for 2 days. After this, gravid females are transferred onto a fresh plate supplemented with yeast paste 1 mM 4 all-trans retinal food. Stage 17 embryos were selected and transferred to the wells coated with dissolved Scotch tape glue, and covered with a layer of halocarbon. Embryos were first recorded for 2 min under infrared light (Andoer IR49S Mini), and then exposed for 2 min with red light or green light and back on infrared light. Videos were acquired at 30 fps and analyzed using a combination of Fiji (ImageJ) and custom-made software LIMAS.

Larvae: Gravid females are transferred onto a fresh plate supplemented with yeast paste 1 mM 4 all-trans retinal food for 24h. Individual freshly hatched larvae were transferred to a new plate with yeast paste 1 mM 4 all-trans retinal food for 12 h and next placed on 0.9% agarose plates for experiments as described above. Crawling larva was first recorded for 30s under infrared light (Andoer IR49S Mini), followed by 30s of red light or green light exposure, and then back on infrared light. The LEDs were controlled with an Arduino ATMega238p 16 MHz (GoTronic) running custom-made GUI software (https://doi.org/10.5281/zenodo.20074803), which itself was controlled through a Raspberry Pi 5 8GB (GoTronic).

Adults: Two-three days individual flies were reared in the dark in the vials containing retinal food (regular fly food mixed with 1mM all-trans retinal (Sigma)) for 5 days. An individual adult fly was placed into a 3D-printed acrylic arena (2×2×0.5cm) with sloped edges^132^, covered by a transparent plexiglass (1mm thick). Fly walking was first recorded for 30s under infrared light (Andoer IR49S Mini), followed by 30s of red light or green light exposure, and then back on infrared light. The LEDs were controlled with an Arduino ATMega238p 16 MHz (GoTronic) running custom-made GUI software (https://doi.org/10.5281/zenodo.20074803), which itself was controlled through a Raspberry Pi 5 8GB (GoTronic).

For all optogenetic experiments, an 8×8 RGB array made up of 64 WS2812E RGB LEDs (AZDelivery) was used and controlled by an Arduino microcontroller with custom-made software to illuminate the arena positioned 3 cm above it. The illumination provided to the samples was 620–630 nm (red light) for CsChrimson and 515–530 nm (green light) for GtACR1.

#### DeepLabCut training and analysis

We employed DeepLabCut (DLC)^35^ to track various body parts from larval and adult locomotion videographies for quantifying particular aspects of this behavior. Next, we first manually labeled three parts of the body (head - thorax and abdomen/tail) for network training and tracking using 200 image frames extracted from distinct videographies. Second, we trained the larval or adult network using a ResNet-50–based neural network for 1,000,000 iterations. We next evaluated the network performance and created a labeled video. We used the labeled videos to check the annotation errors occurred. If these happened, we retrained to improve accuracy with additional labeled 200 images from different videos. If the evaluation of the network is satisfactory, labeled videos are plotted to generate pose estimation data as CSV files, which was analyzed using a custom-made Python code suitable for each developmental stage organism.

### Imaging experiments

#### Immunohistochemistry

Brains and VNCs were dissected in 1 x phosphate-buffered saline (PBS) at distinct time points: 36h ALH for larvae, 90-96h APF pupae, 7-10 days for adults. The tissues were Immunostained as described previously^26^. Briefly, the tissues were then fixed for 1 h in 4% formaldehyde in 1× PBS at room temperature. Of note is the case of embryonic preparations. The embryonic central nervous system (CNS) was dissected to facilitate immunolabelling analysis, as previously described^39^. Embryos were staged for approximately 22 hours at 25°C, and dechorionated using double-sided adhesive tape. The CNS is small, sticky, and difficult to isolate from the body wall muscles. To expose the brain and VNC to immunochemical agents, the body wall was carefully separated using a sharp, thin metal wire (Insect pins, 0.1 mm; Agar Scientific) in 1×PBS. The tissues were then transferred into fixative solution (4% formaldehyde in 1×PBS) and incubated for 20 minutes with gentle shaking. Samples were subsequently washed and incubated with antibody solutions. During the final secondary antibody wash, the dissection was concluded by removing the mouth hooks and any remaining body wall tissue attached to the CNS.

After fixation, CNSs were washed three times (30 min per washing) in PBS with 0.3% Triton X-100 (PBTx) and incubated at 4°C overnight with primary antibodies. The following primary antibodies were used: chicken anti-GFP (Abcam Probes, 1:1000), rat anti-N-cadherin (DSHB DN-Ex, 1:400), mouse monoclonal anti-Antp (DSHB 8C11, 1:100), anti-Pdm2 (Abcam ab201325, 1:200). The secondary antibodies were anti-mouse Alexa Fluor 555 (Invitrogen Molecular Probes, 1:1,000), anti-rabbit Alexa Fluor 647 (Invitrogen Molecular Probes, 1:1,000), and anti-chicken Alexa Fluor 488 (Invitrogen Molecular Probes, 1:1,000). DAPI from ThermoFisher Scientific (D1306) was used to outline the CNS. Images were acquired with a Nikon confocal microscope, processed, and analysed using FIJI ImageJ (NIH).

Neuronal arborization density was quantified by measuring GFP fluorescence in neurites within the VNC prothoracic, mesothoracic, and metathoracic neuromeres. Fluorescence intensity was expressed as corrected total cell fluorescence (CTCF), as previously described^133^. CTCF was calculated as integrated density minus (ROI area × mean background fluorescence). Background fluorescence was determined from regions lacking signal within the same image. *In vivo brain imaging*

Mounting and brain surgery: To prepare flies for imaging, we mounted and performed surgery based on previous methods^134,135^. The flies were first anesthetized on ice (1 min) and transferred to a 4°C aluminium thermoelectric cooling block. Using a multi-axis stage, a pin attached to a custom acrylic holder was coated with UV-curing epoxy (NOA 89, Norland Inc) and lowered onto the fly’s posterior thorax, then cured for 5 s (365/385 nm UV) using a UV wand (Norland Inc). Following a 1 min recovery, we removed the bristles on the fly head using Dumont #5 forceps a dissection microscope (MZ12, Leica) and used a sharp tungsten needle (Tip diameter 1 µm, Fine Science Tools) to perforate the cuticle’s perimeter. This perforated cuticle was then carefully peeled back to expose the brain, and superficial air sacs or fat tissues were gently removed. Ambient humidity was maintained above 40% using a desktop humidifier (Pure Inc). Finally, the open window was sealed with UV-curing epoxy (NOA 68, Norland Inc) and cured for 5 s. For experiments utilizing a water-immersion objective, a #0 cover glass (Electron Microscopy Sciences) was subsequently affixed over the sealed window using NOA 68.

Two-photon imaging in walking adult flies: To track the locomotion of individual flies walking on a trackball during imaging, we used a setup similar to that of prior studies^136,137^. After positioned on the trackball by a three-dimensional translation stage, the fly can navigate on a trackball with two degrees of freedom (forward/backward and left/right rotation). One camera (DFK 33UX290, Imaging Source Inc) records trackball rotation at 314 frame per second, and then we used the FicTrac algorithm^138^ to extract the walking speed and trajectory.

*In vivo* functional calcium imaging was performed using a Thorlabs Bergamo II two-photon microscope with Coherent Axon laser. The jGCaMP8s indicator was excited with 920 nm light from the Axon 920 TPC laser (Coherent Inc). The emitted fluorescence signals were acquired with an Olympus 20X1.0 NA objective (XLUMPLFLN20XW) and through the emission filter cantered at 525 nm with a 50 nm bandwidth at 384 pixels × 256 pixels resolution at 10 Hz imaging rate. The onset of two-photon imaging and behavior recording are synchronized through triggering pulse that was delivered through a multifunction DAQ device (USB-6001, National Instrument) All imaging experiments were conducted at room temperature (∼ 23°C). Behavior and imaging data analysis: Image analysis was performed through custom MATLAB scripts. First, motion correction was applied with the non-rigid ‘NormCorre’ algorithm^139^. Regions of Interest (ROIs) corresponding to the axon branches were then manually selected for each imaging trial. To calculate the *ΔF/F*, the baseline fluorescence (*F*) was defined as the 10th percentile of the fluorescence intensity trace for each imaging trial.

The fly’s locomotion on the ball was quantified using FicTrac to extract forward, angular, and lateral velocities. We defined the fly’s net speed as the root sum squared of these three velocities, as described in prior study^140^, and normalized this value to the maximum speed observed for each individual fly. Moving bouts were defined as periods of sustained locomotion (lasting > 2 s) with an average normalized net speed > 0.15 a.u., which immediately followed a rest period (lasting >1s) with an average normalized speed < 0.05 a.u.

### RT-qPCR

The efficiency of the RNAis used in this study and the expression of dopamine synthesis or transport pathway genes was validated by quantitative PCR with reverse transcription (RT–qPCR). The brain and VNS of female and male flies carrying RNAis against specific gene targets in neurons were dissected and the nSyb-Gal4 pan-neuronal driver were crossed with specific UAS-RNAi or with w1118. RNA was isolated using a standard phenol–chloroform extraction protocol. Residual genomic DNA was removed by DNase treatment for 30 minutes (eZDNAse, Invitrogen). First-strand cDNA synthesis was performed using SuperScript IV First-Strand Synthesis System (Invitrogen), following the manufacturer’s instructions. Quantitative PCR (qPCR) reactions were prepared in a total volume of 10 µL, consisting of 5 µL 2× SYBR Green master mix (LightCycler 480 SYBR Green I Master, Roche), 2 µL nuclease-free water, 1 µL forward primer (5 µM), 1 µL reverse primer (5 µM), and 1 µL cDNA. All samples were run in technical triplicates. Amplification was performed on a LightCycler 480 under the following cycling conditions: denaturation at 95°C for 10 minutes, 35 cycles of 95°C for 15 s, 58°C for 15 s, and 72°C for 10 s, with fluorescence acquisition at the annealing and extension steps. Actin was used as the reference gene. Relative changes in RNA levels of target genes were quantified by normalizing transcript abundance to Actin levels using the 2^(−ΔΔCT)^ method^141^. Primers were designed using FlyPrimerBank^142^ and are listed in the Supplemental Table (**Table S2**).

### Quantification and statistical analysis

Statistical analyses were performed using GraphPad Prism. Data sets were tested for normality using the D’Agostino & Pearson, Anderson-Darling Shapiro-Wilk and Kolmogorov-Smirnov tests. Normally distributed data were tested using one-way ANOVA with Brown–Forsythe and Welch corrections, followed by Dunnett’s T3 multiple-comparisons test. Non-normally distributed data were analyzed using Mann Whitney tests for comparisons between experimental genotypes and their respective parental controls. Friedman tests followed by Dunn’s multiple-comparisons tests were used for within-genotype comparisons across conditions in optogenetic assays. Kruskal–Wallis tests were used for comparisons across conditions in thermogenetic and qPCR assays. Error bars represent the standard error of the mean (SEM). The statistical significance was when p values were <0.05, and is represented as not significant (NS), ∗p < 0.05, ∗∗p < 0.01, ∗∗∗p < 0.001, and ∗∗∗∗p < 0.0001. For each figure panel, the statistical test, error-bar definition, and sample sizes are indicated in the legends.

## Supplemental information

**Figure S1.**
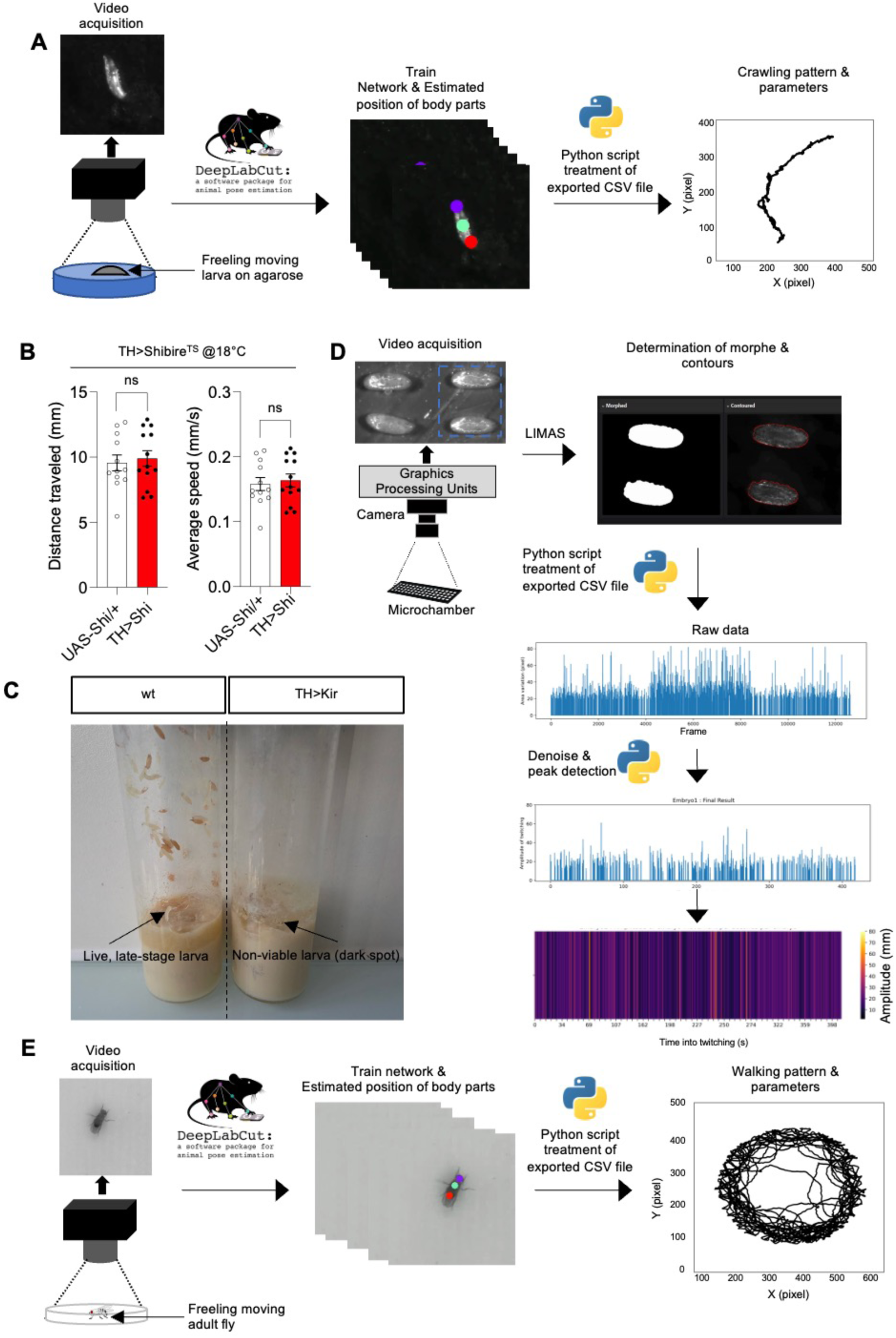
Suppressing DAN activity during embryogenesis disrupts locomotor development, related to Figure 1. **A.** Protocol to monitor larval locomotion. We applied DeepLabCut along with a custom-made Python script to analyze larval crawling patterns from video recordings. **B.** At the permissive temperature of 18°C, the expression of Shibire^TS^ (Shi) in all DANs (TH>Shi) does not affect larval crawling components, distance traveled and average speed, compared to the control (*UAS-Shi/+*). Mean ± SEM. Error bars: 95% CI. ns p >0.05 by Mann-Whitney test. n for *UAS-Shi/+ and TH>Shi*: 12 and 13, respectively. Data were collected from two independent experiments, and each animal was recorded individually. **C.** Representative photographs showing impaired wandering behavior in larvae expressing Kir in DANs (*TH>Kir*). **D.** Protocol to monitor embryonic locomotor activity. We used a custom-developed software, Lifelong Movement Analysis Software (LIMAS), along with a Python script to detect embryonic twitching. **E.** Protocol to monitor adult fly locomotion. We applied DeepLabCut^35^ and a custom-made Python script to analyze adult fly walking patterns from videos.

**Figure S2.**
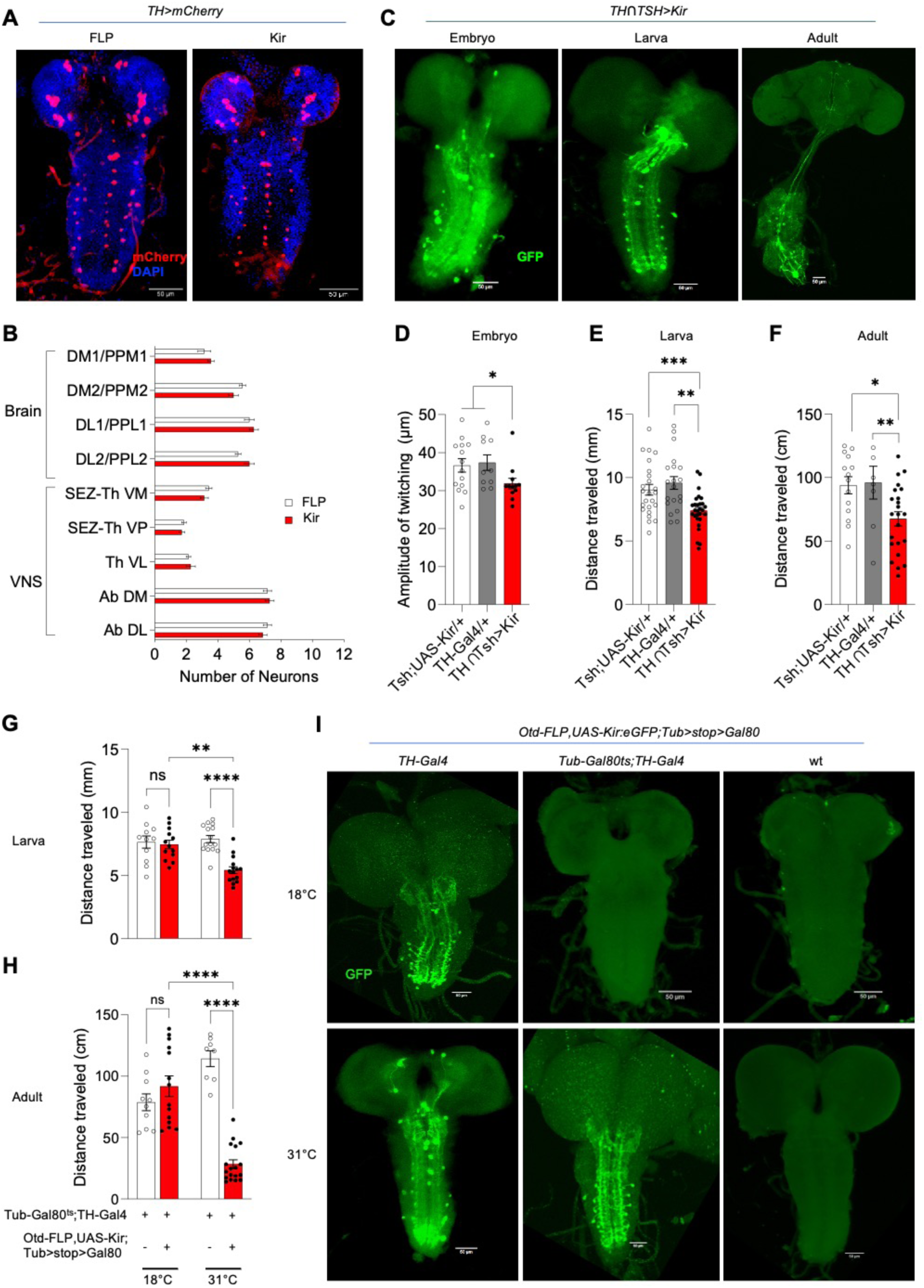
Early activity of VNS DANs is required for both early and adult locomotor function, related to Figure 2. **A, B.** Confocal images (**A**) show that Kir expression in dopamine neurons (TH>Kir) does not affect their cell number (**B**). n=7. **C–F.** Constitutive suppression of activity in VNS DANs alters locomotion during development and adulthood. Confocal images showing Kir-GFP expression in VNS DANs across distinct developmental stages (C). We note a loss of neurite arborization, which may be related to the Kir-mediated electrical activity suppression effect. Embryonic twitching amplitude (D), as well as larval (E) and adult (F) traveled distance, are reduced compared to controls. Mean ± SEM. Error bars: 95% CI. ∗p < 0.05, ∗∗p < 0.01 and ∗∗∗p < 0.001 by Mann-Whitney test. n for UAS-Kir/+, TH-Gal4/+, and TH∩TSH>Kir were 14, 10, and 12 (embryo); 24, 20, and 28 (larva); and 14, 8, and 23 (adult), respectively. Data were collected from two independent experiments, and each animal was recorded individually. **G, H.** Long-lasting locomotor effects following temporal suppression of activity in VNS DANs during the embryonic stage. Larvae (G) and adults (H) that have experience Kir expression at embryonic stage exhibit short distance traveled. Mean ± SEM. Error bars: 95% CI. ns p >0.05, ∗∗p < 0.01, and ∗∗∗∗p < 0.0001 by Kruskal-Wallis test with Dunn’s multiple comparisons test. n for tub-Gal80^ts^;*TH-Gal4/+* and *Otd-FLP/UAS-Kir,Tub-Gal80^ts^;TH-Gal4/Tub>stop>Gal80* at 18°C were 11 and 13 (larva); 10 and 14 (adult), respectively. n for tub-Gal80^ts^;*TH-Gal4/+ and Otd-FLP/UAS-Kir,Tub-Gal80^ts^;TH-Gal4/Tub>stop>Gal80* at 31°C: 15 and 15 (larva), 8 and 18 (adult) respectively. Each animal was recorded individually. **I**. Verification of Tub-Gal80^ts^ approach efficiency. Confocal images showing Kir-GFP expression in the VNS of larvae carrying the TH-Gal4 driver, which drives the expression in DANs. Larvae were raised from embryonic stage at 18°C or 31°C. At 18°C, larvae expressing Gal80^ts^ showed undetectable GFP fluorescence in the VNS, whereas at 31°C, Kir-GFP expression was detected in VNS DANs (middle). Control larvae with (left) or without TH-Gal4 (right) showed GFP expression or no fluorescence, respectively.

**Figure S3.**
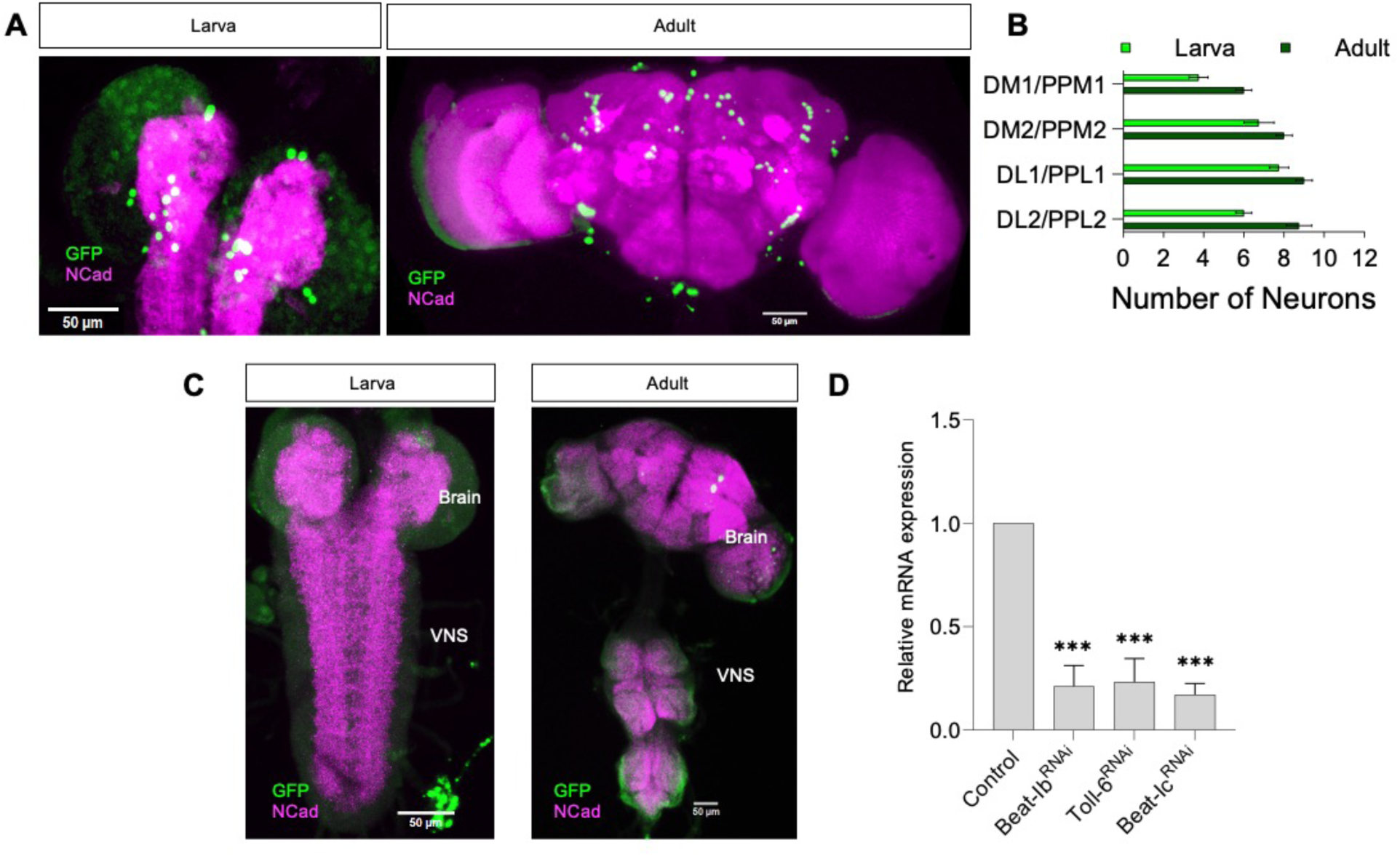
DANs across development, related to Figure 3 and 4**. A-C**. Immortalization of a DANs. Confocal images (left) and quantification (right) of the spatial distribution of nEGFP^+^ DANs in larval and adult brain reveals no difference in cell numbers (B). No GFP^+^ neurons in the VNC and brain of controls flies reared at 18°C (C). n=6-7 individual animals. **D.** mRNA levels of Beat-Ib, Toll-6, and Beat-Ic following RNAi expression. RT–qPCR was performed to assess the efficiency of different RNAi constructs in the adult nervous system following pan-neuronal expression, using the 2^(-ΔΔCT)^ method. Relative mRNA expression levels were normalized to β-Actin transcripts as an internal control and expressed relative to control samples set to 1. RNAi-mediated knockdown of Beat-Ib, Toll-6, and Beat-Ic resulted in reduced mRNA levels compared to control (nSyb>GFP). Mean ± SEM. ∗p < 0.05, and ∗∗p < 0.01 by Kruskal–Wallis test followed by Dunn’s multiple comparisons test. Analyses were performed using three independent RT-qPCR experiments carried out on three different RNA extractions from ten to twelve male and female nervous systems per genotype.

**Figure S4.**
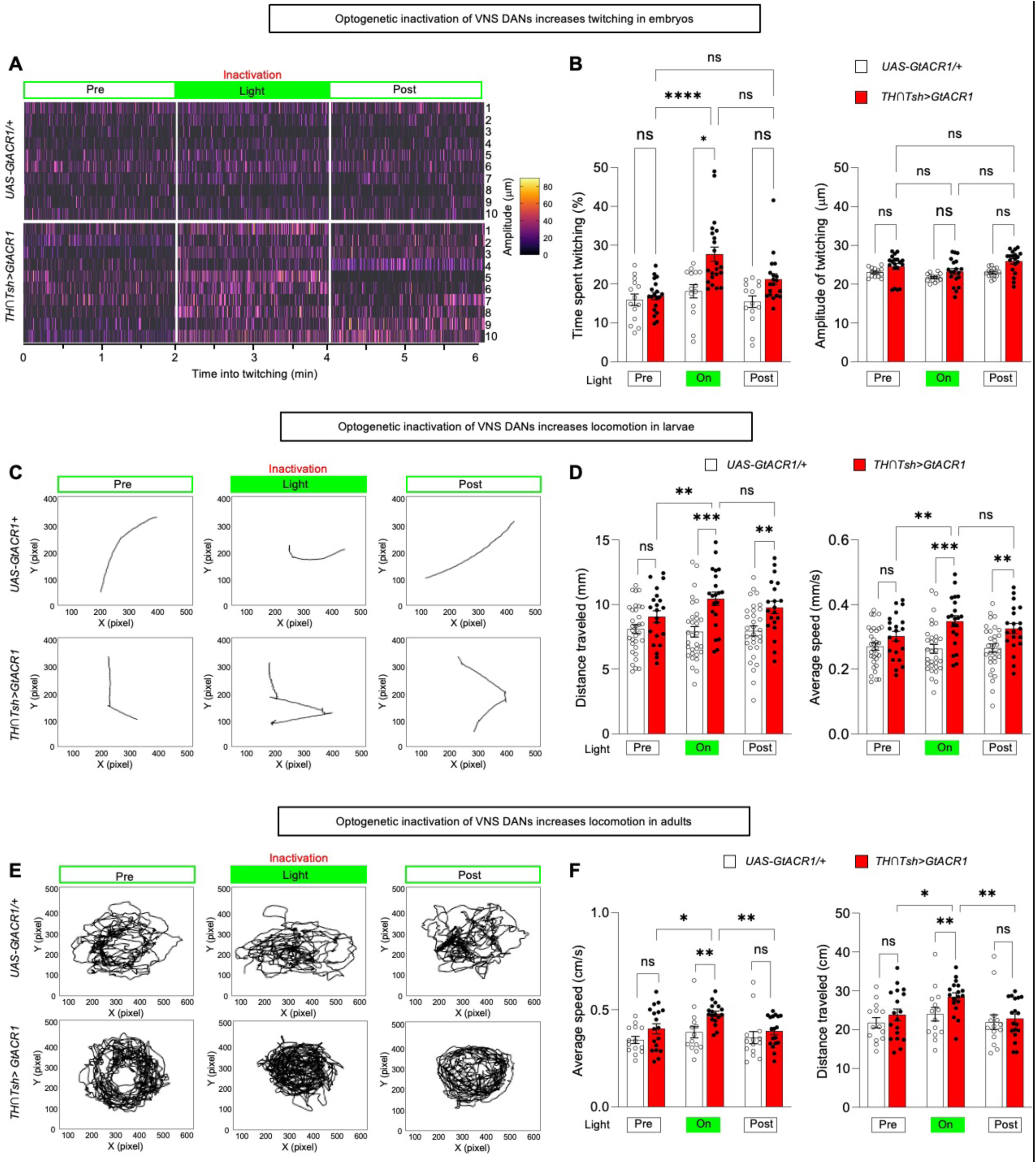
Acute inactivation of VNS DANs reduces locomotor speed across distinct developmental stage, related to Figure 5. **A, B**. Embryonic twitching. Heatmaps of embryos show that green-light stimulation increases twitching in embryos expressing GtACR1 in VNC DANs (TH∩Tsh>GtACR1), compared to pre-and post-light periods and genetic controls (UAS-GtACR1/+) (A). Heat hatches denote twitching and each row represent individual embryo. Quantification of time spent twitching and twitching amplitude (B). Mean ± SEM. Error bars: 95% CI. ∗p < 0.05, ∗∗p < 0.01, and ∗∗∗p < 0.001 by Mann Whitney tests for comparisons with parental controls; Friedman tests followed by Dunn’s tests for within-genotype comparisons. n for *UAS-GtACR1*/+ and TH∩Tsh>GtACR1 for all periods: 14 and 19, respectively. Data were collected from two independent experiments, and each animal was recorded individually. **C, D**. Larval locomotion. Representative crawling patterns show longer crawling paths in TH∩Tsh>GtACR1 larvae during green-light stimulation, compared to pre-and post-light periods and controls (*UAS-GtACR1*/+) (C). Quantification of distance traveled and crawling speed (D). Mean ± SEM. Error bars: 95% CI. ∗p < 0.05, ∗∗p < 0.01, and ∗∗∗p < 0.001 by Mann Whitney tests for comparisons with parental controls; Friedman tests followed by Dunn’s tests for within-genotype comparisons. n for UAS-GtACR1/+ and TH∩Tsh>GtACR1 for all periods: 23 and 17, respectively. Each animal was recorded individually across two independent experiments. **E, F**. Adult locomotion. Representative patterns show longer walking paths in *TH∩Tsh>GtACR1* adults during green-light stimulation, compared to pre-and post-light periods and controls (*UAS-GtACR1*/+) (E). Quantification of distance traveled and walking speed (F). Mean ± SEM. Error bars: 95% CI. ∗p < 0.05, ∗∗p < 0.01, and ∗∗∗p < 0.001 by Mann Whitney tests for comparisons with parental controls; Friedman tests followed by Dunn’s tests for within-genotype comparisons. n for *UAS-GtACR1*/+ and *TH∩Tsh>GtACR1* for all periods: 14 and 20, respectively. Data were collected from two independent experiments, and each animal was recorded individually.

**Figure S5.**
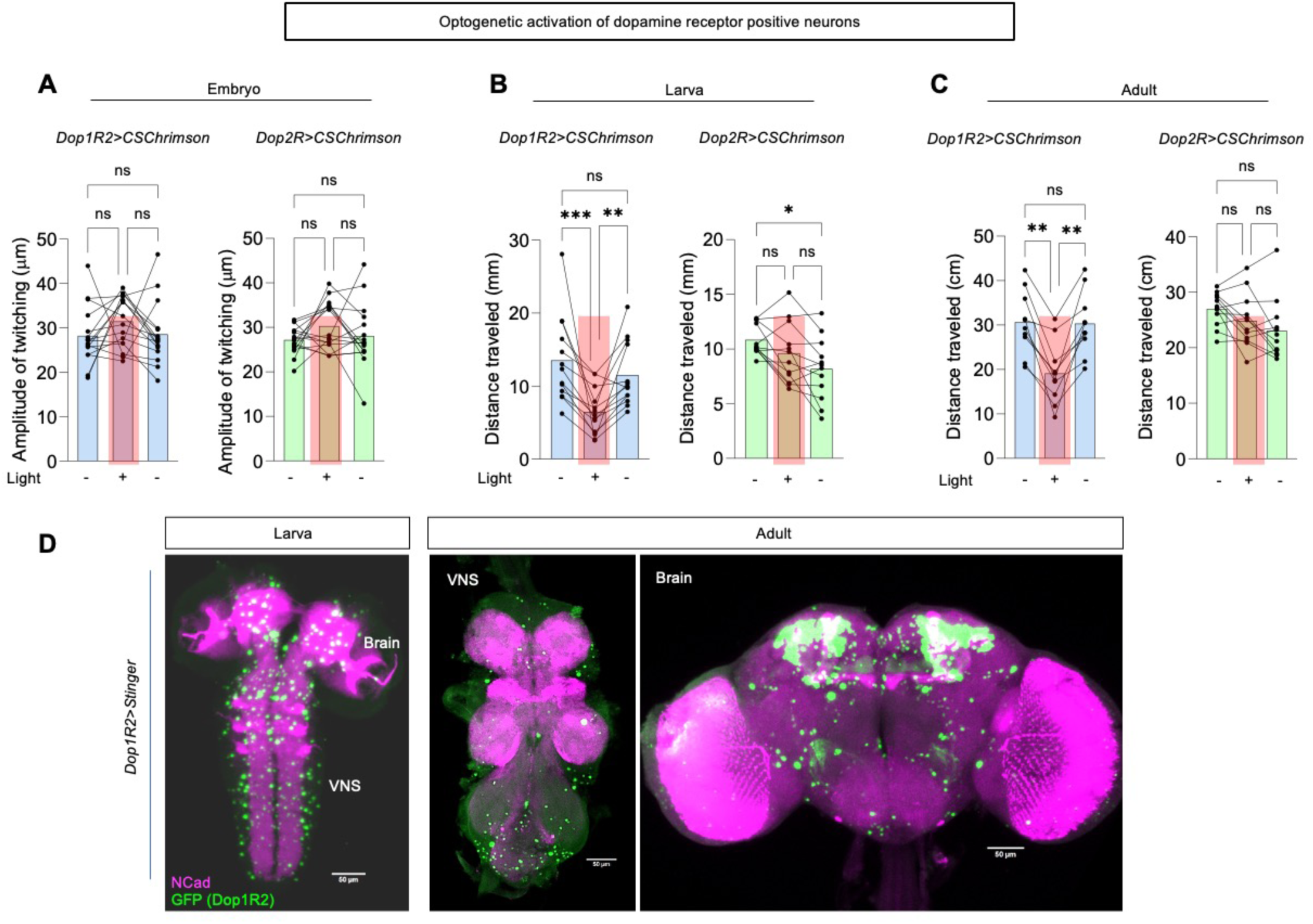
Acute activity of dopamine receptors and locomotor performance, related to Figure 6. **A–C**. Red-light stimulation effects on embryonic twitching amplitude (A), larval crawling (B), and adult walking (C) in animals expressing Chrimson::mVenus in Dop1R2^+^ neurons (*Dop1R2>CsChrimson*) (left), compared to pre-and post-light periods and *Dop2R>CsChrimson* flies (right). The stimulation has no effect on embryonic twitching amplitude but reduces larval and adult traveled distance. Mean ± SEM. ns p>0.05, ∗p < 0.05, ∗∗p < 0.01 and ∗∗∗p < 0.001 by one-way ANOVA with Dunnett’s T3 multiple comparisons test. n for *Dop1R2>CsChrimson* and *Dop2R>CsChrimson* for all periods: 16 and 16 (embryo), 13 and 12 (larva), and 10 and 12 (adult). Data were collected from two independent experiments, and each animal was recorded individually. **D**. Confocal images of Dop1R2⁺ neurons expressing nuclear GFP (Dop1R2>stinger) in the larval and adult nervous system. They show that Dop1R2⁺ neurons (green dots) are distributed in both the brain and the VNS.

**Figure S6.**
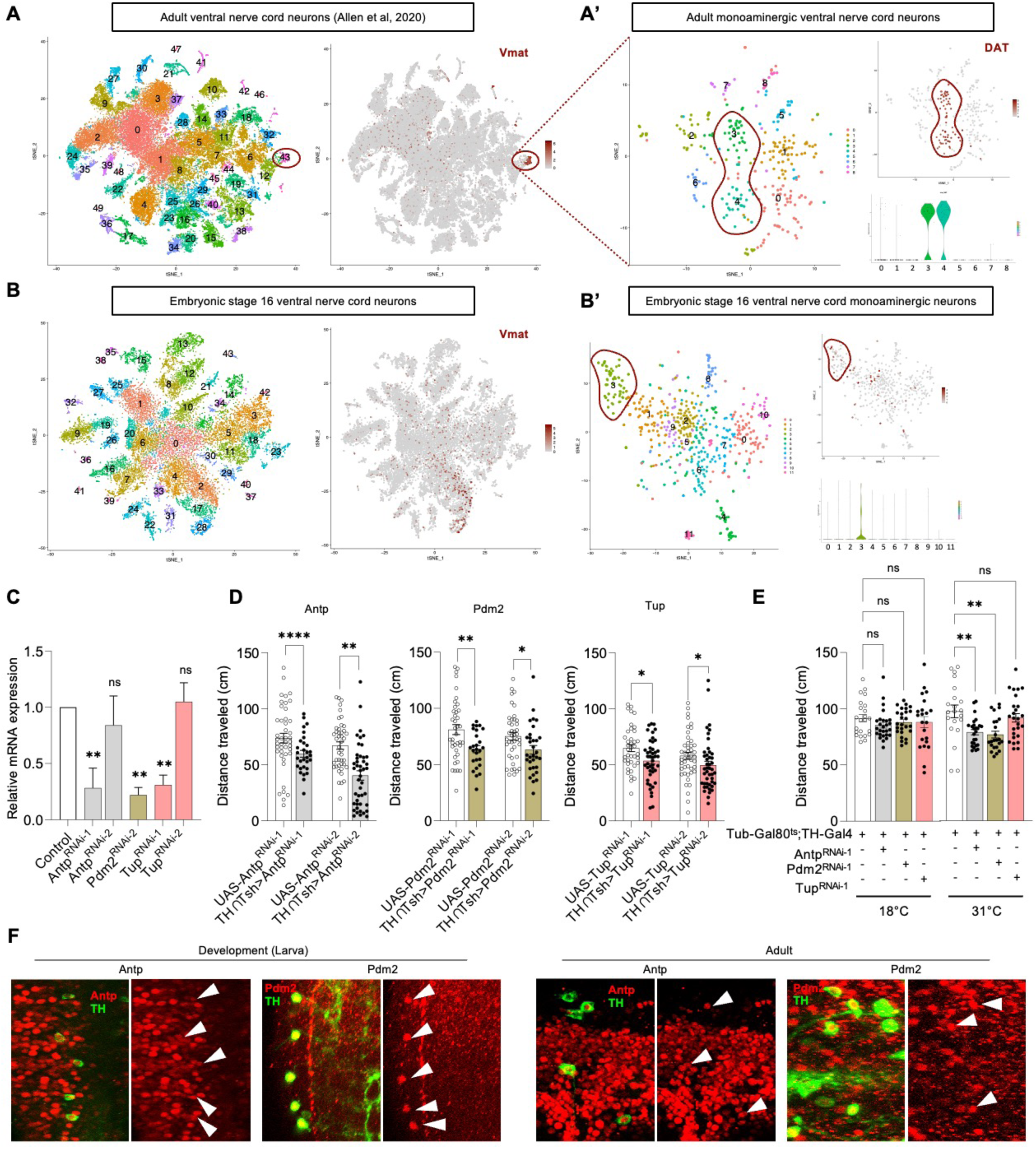
Developmental genetic programs are reused by VNS DANs to sustain locomotor function, related to Figure 7. **A.** tSNE plot of the all adult ventral nerve cord neurons (as sequenced by Allen et al, 2020^22^). Cluster 43 is enriched in the expression of Vmat, a marker for monoaminergic neurons. The adult monoaminergic ventral nerve cord neurons (i.e. cluster 43) were re-analysed and re-clustered. Clusters 3 and 4 are enriched in the expression of DAT, marker of dopaminergic neurons, as shown by a feature plot and a violin plot (A’). **B.** tSNE plot of the all stage 16 ventral nerve cord neurons (as sequenced by Seroka et al, 2022^68^). Monoaminergic neurons can be found throughout and do not form a specific cluster. Vmat-positive cells were re-analysed and re-clustered. Cluster 3 is enriched in the expression of DAT, marker of DANs, as shown by a feature plot and a violin plot (B’). **C.** mRNA levels of Antp, Pdm2, and Tup following RNAi expression. RT–qPCR was performed to assess the efficiency of different RNAi constructs in the adult nervous system following pan-neuronal expression, using the 2^(-ΔΔCT)^ method. Relative mRNA expression levels were normalized to β-Actin transcripts as an internal control and expressed relative to control samples set to 1. The expression of Antp^RNAi-1^ (VDRC_KK101774), Pdm2^RNAi-1^ (BDSC_50665), and Tup^RNAi-1^ (VDRC_KK103585) results in reduced mRNA levels compared to control (nSyb>GFP). Pan-neuronal (nSyb-Gal4) expression of Pdm2^RNAi-2^ produced no progeny, likely due to UAS-Pdm2^RNAi-2^ homozygous sterility. ∗p < 0.05, and ∗∗p < 0.01 by Kruskal–Wallis test followed by Dunn’s multiple comparisons test. Analyses were performed using three independent RT-qPCR experiments carried out on three different RNA extractions from ten to twelve male and female nervous systems per genotype. **D.** Constitutive RNAi-mediated downregulation of selected developmental transcription factors, including Antp, Pdm2, and Tup, reduces the distance traveled by adult flies. Mean ± SEM. Error bars: 95% CI. ∗p < 0.05, ∗∗p < 0.01, ∗∗∗p < 0.001 and ∗∗∗∗p < 0.0001 by Mann-Whitney tests. n for *UAS-Antp^RNAi-1^*, *TH∩Tsh>Antp^RNAi-1^*, *UAS-Antp^RNAi-2^*, *TH∩Tsh>Antp^RNAi-2^*, *UAS-Pdm2^RNAi-1^*, *TH∩Tsh>Pdm2^RNAi-1^*, *UAS-Pdm2^RNAi-2^*, *TH∩Tsh>Pdm2^RNAi-2^*, *UAS-Tup^RNAi-1^*, *TH∩Tsh>Tup^RNAi-1^*, *UAS-Tup^RNAi-2^* and *TH∩Tsh>Tup^RNAi-2^*: 45, 30, 44, 45, 39, 26, 42, 32, 41, 45, 45, and 45. Data were collected from three independent experiments, and each animal was recorded individually. **E.** Adult-specific downregulation of Antp andPdm2, but not Tup, in DANs reduced fly distance travelled, suggesting an impaired locomotion. Mean ± SEM. ns p >0.05 and ∗∗p < 0.01 by Kruskal-Wallis test with Dunn’s multiple comparisons test. n for *Tub-Gal80^ts^;TH-Gal4/+, Antp^RNAi-1^, Pdm2^RNAi-1^ and Tup^RNAi-1^* transgenic lines: 23, 27, 26, and 22 at 18°C, and 22, 21, 26, and 23 at 31°C, respectively. Data were collected from three independent experiments, and each animal was recorded individually. **F.** Verification of Antp and Pdm2 protein expression in DANs during development (left) and adulthood (right). All scale bars are 10 μm.

